# A combination TLR7/8 and RIG-I agonist adjuvant reverts asthmatic allergic sensitization and prevents aggravated influenza infection in OVA-sensitized mice

**DOI:** 10.1101/2025.06.23.659362

**Authors:** Juan García-Bernalt Diego, Eleanor Burgess, Lauren A. Chang, Matthew Prellberg, Moataz Noureddine, Naseem Sadek, Yong Chen, Farah El-ayache, Gabriel Laghlali, Seok-Chan Park, Vivian Yan, Pamela T Wong, Bruno G. De Geest, Jeffrey A. Tomalka, Michael Schotsaert

## Abstract

Allergen-specific immunotherapy (AIT) is the only disease-modifying treatment currently available to treat allergy. However, it has limitations, as most allergens are poorly immunogenic, resulting in an AIT process that can take years. Therefore, adjuvant selection becomes critical to achieve a more efficacious therapy. Our group has developed and tested an amphiphilic TLR7/8 agonist (IMDQ) and a RIG-I agonist (SDI) that used alone, or in combination, have demonstrated strong adjuvant activity for influenza and SARS-CoV-2 vaccines in preclinical models. Here we describe the effect of these adjuvants in the sensitization of preclinical models with the ovalbumin (OVA) asthmatic allergic model via an in-depth humoral and cellular immune profiling. We assess their immune skewing and tolerance inducing capacities in previously sensitized preclinical models with different genetic backgrounds (C57BL/6 *vs.* BALB/c mice). Moreover, we evaluate their effect in an unrelated antigenic challenge with influenza. Finally, we investigate the role of IgG subclasses and T-cell subpopulations in the protection against OVA challenge conferred by the combination of IMDQ and SDI. We demonstrate that OVA-immunization in combination with IMDQ+SDI prevents allergic sensitization via the induction of a balanced Type 1/Type 2 response. Furthermore, it can revert the allergic phenotype in mice previously sensitized with OVA-Alum, through reducing lung eosinophilia, as well as IL-4 and IL-5 production. However, this was dependent on genetic background. IMDQ+SDI sensitization also led to reduced morbidity of a secondary influenza challenge in OVA-sensitized mice. Finally, we demonstrated that IgG2c, by itself, cannot protect from allergic sensitization and that both CD4^+^ and CD8^+^ T-cells are needed for IMDQ+SDI prevention of eosinophil recruitment and activation upon intranasal OVA-challenge.

## INTRODUCTION

Asthmatic allergy is defined as a heterogeneous disease characterized by chronic airway inflammation and remodeling. Its origin and severity are strongly determined by genetic and environmental factors, and in most cases, associated with a childhood IgE-dependent sensitization to environmental allergens, although it can also emerge later in life^1^. Based on data from the 2019 Global Burden of Diseases, Injuries, and Risk Factors Study (GBD), asthma prevalence continues to rise, with an estimated burden of 262.41 million cases globally^2^. Besides its obvious relevance as a primary cause of morbidity, it has been identified as a comorbidity for numerous conditions including rhinitis, sinusitis, gastroesophageal reflux disease or obstructive sleep apnea^3^. Additionally, some respiratory infections, such as influenza or invasive pneumococcal disease, have been associated with asthma exacerbations or enhanced pathology of the infection in asthmatic patients^4,5^ Currently, the only disease-modifying treatment for allergies is allergen-specific immunotherapy (AIT). To perform this treatment, patients are confronted with increasing amounts of the allergen over long periods of time. If successful, AIT can result in tolerance towards the offending allergen and can lead to a late decline in allergen-specific IgE, together with an early increase in allergen-specific IgG subclasses and the induction of regulatory T-and B-cell subsets^6,7^. While this is a sound strategy, it has limitations, as pure forms of most allergens are poorly immunogenic, resulting in an AIT process that can take up to three years. Thus, appropriate adjuvantation leading to more efficacious therapy with faster onset of action is of the utmost importance to improve AIT^8,9^.

An ideal adjuvant, when combined with an allergy vaccine, should activate the innate immune system to amplify the downstream adaptive response to the antigen. Additionally, it should be biodegradable, stable, sustainable, nontoxic, and cost-effective. As of 2020, only four adjuvants have been approved for commercial use in AIT: aluminum hydroxide, calcium phosphate, microcrystalline tyrosine (MCT) and monophosphoryl lipid A (MPL)^10^. Aluminum salts (from now on referred to as Alum) have been used as an adjuvant for almost a century. While it is routinely used in AIT because it can improve allergen immunogenicity and tolerability, it can also boost undesired IgE and lead to a Type 2 immune response polarization^11^. Notwithstanding, these clinically undesired features allow the use of this adjuvant to sensitized preclinical models for allergic asthma induction^12^.

In contrast to adjuvants such as Alum, Toll-like receptor (TLR) agonists and RIG-I agonists have a clear immunomodulatory profile that could favor anti-allergic T-cell responses. This could be a promising avenue for AIT improvement. The anti-allergic properties of TLRs are characterized by a Type 1 immune skewing, induction of regulatory responses, and production of blocking antibodies. Nonetheless, different TLR agonists are in very distant stages in their development towards AIT. Synthetic TLR4 and TLR9 agonists have already entered clinical trials, while TLR2, TLR5 and TLR7/8 agonists shown a potent anti-allergic effect in *in-vitro* and preclinical models but have yet to reached human trial^9^. While RIG-I agonists are known to induce Type 1 polarization and T-cell response induction through IRF3^13^ and they are under intense study in viral vaccinology^14,15^ as well as cancer immunotherapy^16^, their implementation in AIT is absent.

Our group has developed and tested an amphiphilic TLR7/8 agonist based on an imidazoquinoline that is conjugated to a 3 kDa poly(ethylene glycol) (PEG) with a cholesteryl motif at the opposite chain end of the PEG. This compound IMDQ-PEG-Chol, (referred to hereafter as IMDQ), has shown its immunopotentiating and immune-polarizing capabilities in combination with both influenza^14^ and SARS-CoV-2 vaccines^17^. Additionally, we have produced a Sendai virus-derived RNA agonist of RIG-I (referred to hereafter as SDI) and proved its potential as an influenza vaccine adjuvant^18^ and SARS-CoV-2 vaccine adjuvant^15^. Moreover, combination of both IMDQ and SDI has shown to completely skew the vaccine-induced influenza immune response towards IgG2a in a BALB/c model, also resulting in an induction of HAI titers and antibody-mediated cellular cytotoxicity (ADCC), as well as improved T-cell responses that correlated with protection against lethal influenza challenge. These adjuvants, either alone or in combination, yielded better results than commercially used AddaVax^14^.

We want to explore if the strong Type 1 immune polarization obtained with these adjuvants in the context of anti-viral vaccines suggest that they might be good candidates for AIT. In this manuscript we describe the effect of these adjuvants, alone and in combination, in the sensitization of preclinical models with the standard ovalbumin (OVA) asthmatic allergic model. For that, we perform an in-depth humoral and cellular immune profiling of the models after sensitization and OVA challenge. Furthermore, we assess their immune skewing and tolerance inducing capacities in previously sensitized mice with different genetic backgrounds (C57BL/6 and BALB/c mice). Additionally, we evaluate their role in the lung remodeling and enhancement of influenza pathology in sensitized mice. Finally, we investigate the role of IgG immunoglobulin subtypes and T-cell subpopulations in the protection against allergy conferred by the combination of IMDQ and SDI.

## RESULTS

### OVA sensitization in combination with IMDQ or IMDQ+SDI leads to a balanced Type 1/Type 2 immune response to intranasal OVA challenge

First, we assessed if IMDQ, alone or in combination with SDI, could polarize the immune response to OVA towards a Type 1 or balanced Type 1/Type 2 response. 6-8 week-old female C57BL/6 (n=46) were immunized intramuscularly twice, one week apart, with 20 µg of OVA in combination with Alum, a Type 2 skewing adjuvant (n=10) or with Type 1 skewing adjuvants such as IMDQ (n=10) or IMDQ+SDI (n=10). An SDI alone experimental group was not included in this study, as previous results have shown that it induces a balanced, slightly Type 2 skewed response in the absence of IMDQ, and thus would not qualify as type 1 skewing adjuvant^14^. An additional group received only PBS intramuscularly as a negative control (n=8) and one group received OVA-Alum intraperitoneally (n=8), the standard sensitization route for asthmatic allergic induction^19^, as a positive control (Fig. 1A). Mice showed detectable OVA-binding IgG as soon as seven days after the first sensitization (-7DPC, days-post challenge), particularly those sensitized with OVA-Alum intraperitoneally [OVA-Alum (IP)], and the OVA-IMDQ+SDI group. Prior to the intranasal OVA challenge, no significant differences were found among sensitized groups. However, one day after the challenge (1DPC), OVA-Alum (IP) showed significantly higher OVA-binding IgG than the rest of the groups. These titers rapidly declined and at 11DPC groups receiving Type 1 skewing adjuvants showed significantly higher OVA-binding IgG compared to OVA-Alum (IP) group. We then focused on IgG subclasses. In C57BL/6, antigen-specific IgG1 and IgG2c are commonly used as serological markers of Th2- and Th1-skewed responses, respectively, as IL-4 promotes class switching to IgG1 and IFN-γ to IgG2c^20^. The Alum groups, showed significantly higher OVA-binding IgG1 titers than IMDQ or IMDQ+SDI groups, even prior to challenge (0DPC). OVA challenge increased those differences which remained significant until the end of the experiment. On the other hand, none of the Alum groups presented detectable OVA-binding IgG2c titers, while this antibody subclass was induced in OVA-IMDQ and OVA-IMDQ+SDI groups as soon as one week after the first sensitization, was boosted after the second sensitization, and was then maintained until the end of the experiment (Fig. 1B). In all, groups sensitized with OVA-Alum, regardless of the administration route, exhibit a highly Type 2 skewed antibody response, while mice sensitized with OVA-IMDQ or OVA-IMDQ+SDI showed a more balance, slightly Type 1 skewed antibody response (Fig 1C).

**Figure 1.**
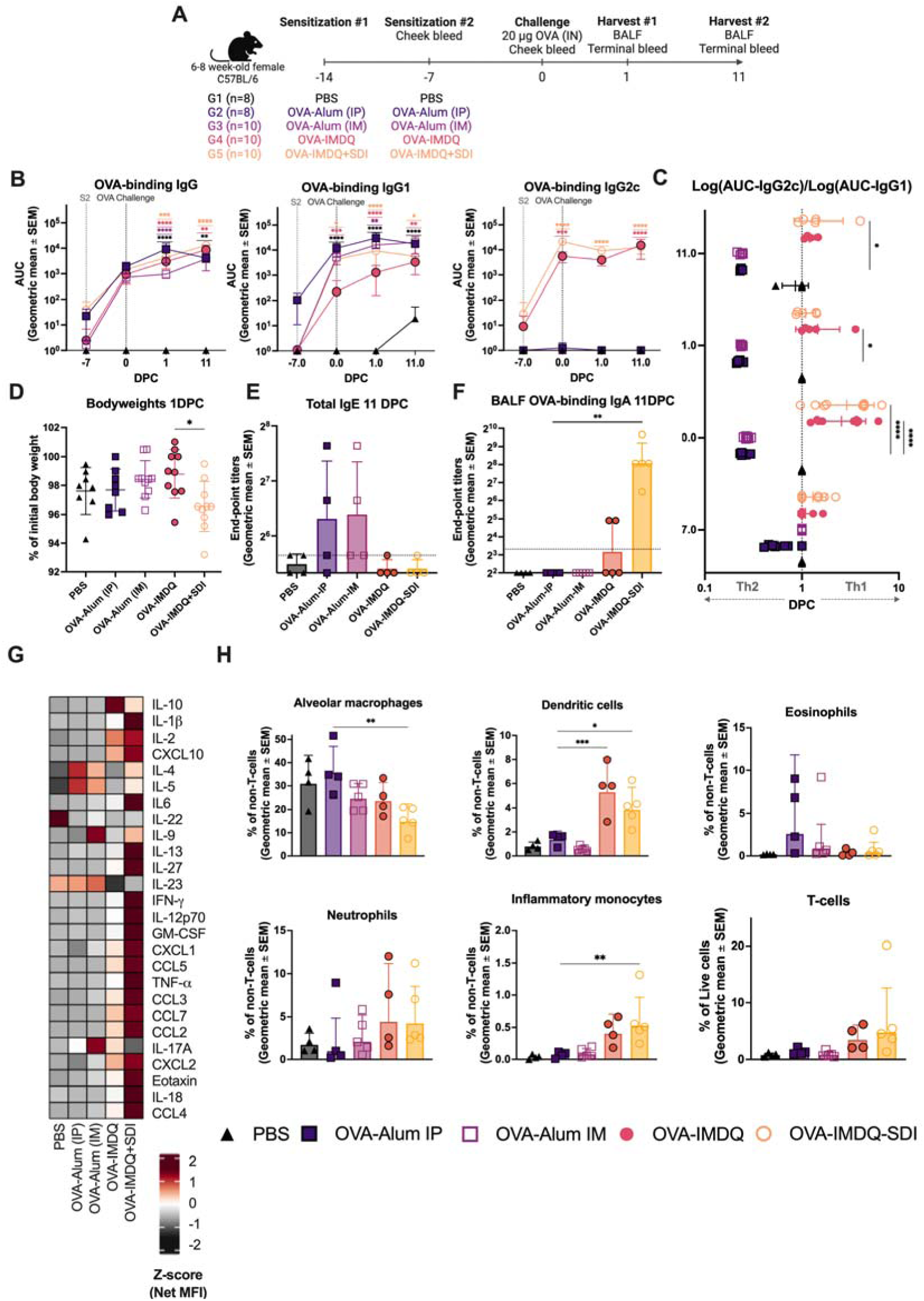
OVA sensitization with Th1-skewing IMDQ and IMDQ-SDI leads to a balanced Th1/Th2 response to the allergen in naïve C57BL/6. A) Experimental design. Created with: <u>Biorender.com</u>. B) From left to right: OVA-binding IgG, OVA-binding IgG1 and OVA-binding IgG2c represented as Area under the curve (AUC) of the optical density at 450 nm (OD450) minus the background noise (OD650). C) Skewing of the antibody response is represented as the coefficient between the log of the OVA-binding IgG2c (AUC) and the log of the OVA-binding IgG1 (AUC). Minimum values are set to 1 for representation purposes. D) Bodyweight loss of mice 1 day after the challenge with OVA. E) End-point titers of Total IgE measured in serum at 11DPC. Limit of detection is established at 50, represented by the dotted line in the graph. F) End-point titers of OVA-binding IgA measured in BALF at 11DPC. Limit of detection is established at 10, represented by the dotted line in the graph. G) Heatmap of net mean fluorescence intensity (MFI) for 26 cytokines/chemokines, z-scored by row, measured in BALF 1DPC. H) Percentage of BALF cell populations measured by flow cytometry 1DPC. From left to right and top to bottom: alveolar macrophages (Gating: Live > CD3^-^ > Ly6G^-^ > CD11c^+^ CD11b^-^ > SiglecF^+^), dendritic cells (Gating: Live > CD3^-^ > Ly6G^-^ > CD11c^+^>MHCII^+^ SiglecF^-^), eosinophils (Gating: Live > CD3^-^ > Ly6G^-^ > CD11c^-^ CD11b^+^ > SiglecF^+^), neutrophils (Gating: Live > CD3^-^ > Ly6G^+^ CD11b^+^), Inflammatory monocytes (Gating: Live>CD3^-^> Ly6G^-^ > CD11c^-^ CD11b^-^ > Ly6C^+^) and T-cells (Gating: Live>CD3^+^). Statistical analysis: for comparisons among groups at several time points: Two-way ANOVA with Tukey’s multiple comparisons test using the OVA-Alum (IP) group as control for multiple comparisons. Color of the significance markers indicates the comparison they represent; for comparisons among groups at a specific time point: Kruskal-Wallis one-way ANOVA with Dunn’s multiple comparisons test using the OVA-Alum (IP) group as control for multiple comparisons. Significance is represented as:*p = 0.05 to 0.01, **p = 0.01 to 0.001, ***p = 0.001 to 0.0001, ***p < 0.0001.

For the OVA intranasal challenge, mice were mildly anesthetized with ketamine/xylazine. In that context, a small drop in body weight 1DPC is expected. Still, the OVA-IMDQ+SDI group lost significantly more weight than OVA-IMDQ (Fig 1D). Although the difference was significant, body weight loss was transient in all groups and initial body weight was recovered by 2DPC and remained stable 7DPC.

Allergic responses are characterized by IgE production, although C57BL/6 mice do not produce as high IgE levels as other mouse strains such as BALB/c^21^. Nonetheless, total IgE was measured in serum at 11DPC. IgE could only be detected in some mice from the OVA-Alum (IP) and OVA-Alum (IM) groups, but not in the rest. Notwithstanding, IgE levels were low, and the increase was not statistically significant (Fig. 1E).

Clinical observations have linked IgA with improved allergic tolerance^22^. To assess mucosal antibody responses, OVA-binding IgA was measured in bronchoalveolar lavage fluid (BALF) at 11DPC. Two of five animals in the OVA-IMDQ group exhibited detectable OVA-binding IgA, while all animals from the OVA-IMDQ+SDI group presented OVA-binding IgA (Fig. 1F). None of the animals in the Alum groups presented detectable OVA-binding IgA.

Cytokine profiling performed by multiplex ELISA and cellular profiling performed by flow cytometry in BALF, supports the immune polarization described in the humoral response. Both OVA-Alum (IP) and OVA-Alum (IM) groups presented elevated levels of the Type 2-cytokines IL-4 and IL-5, as well as IL-23. Additionally, OVA-Alum (IM) groups presented elevated levels of the key Type 2 cytokine IL-9 and as well as IL-17A, typically associated with Type 17 responses and neutrophilic inflammation. At 11DPC, while levels of IL-4 and IL-5 were decreased in both groups, IL-9, IL-17A, IL-22 and IL-23 remained elevated. Those two groups also presented a non-significant increase in the fraction of eosinophils in BALF at 1DPC, which remained elevated until 11DPC, as well as a higher fraction of alveolar macrophages (AMs) only at 1DPC (Fig. 1G, H, S1-3).

On the other hand, OVA-IMDQ and OVA-IMDQ+SDI groups showed a strong proinflammatory Type 1 skewed cytokine/chemokine profile in BALF. The OVA-IMDQ group only showed significant increase in CXCL-10 and MIP-2[when compared to OVA-Alum (IP) (Fig. 1G, S1 and S2). The OVA-IMDQ+SDI group presented a much more extreme profile with significant increases of IL-2, CXCL-10, INF-[, GRO-[, TNF-[, MIP-1[, CCL-7, MIP-2[, IL-18 and MIP-1β (Fig 1G, S1 and S2). Additionally, those groups presented a recruitment of neutrophils, dendritic cells (DCs), inflammatory monocytes (IMs) and T-cells into the BALF, as soon as 1DPC. 11DPC T-cells and dendritic cells remained elevated. At 11DPC, all cytokines were back to baseline levels except for IL-2 in the OVA-IMDQ group which remained significantly elevated when compared to OVA-Alum (IP). Interestingly, OVA-IMDQ+SDI showed significantly higher levels of IL-13 at 1DPC, as well as the eosinophil recruiting chemokine Eotaxin, compared to OVA-Alum (IP), but no increased eosinophil recruitment (Fig. 1G, H, S1-S3).

### IMDQ+SDI prevents allergic response in previously sensitized C57BL/6 and ameliorates it in BALB/c

Once we proved that IMDQ or IMDQ+SDI could skew the immune response to an intranasal OVA-challenge towards a balanced Type 1/Type 2 response in naïve mice, we evaluated the flexibility of the skewing in previously sensitized mice. Thus, 6–8-week-old female C57BL/6 mice (n=48) were divided into four groups that were all sensitized intramuscularly four times, with one week between sensitizations. One received PBS four times, as a negative control and one received OVA-Alum (referred to as Alum onward) intramuscularly four times as a positive control. A third group was injected twice with OVA-Alum to establish an allergy sensitized model as we did in the previous experiment but then received two doses of OVA-IMDQ+SDI prior to challenge (indicated hereafter as Alum➔IMDQ+SDI). A final group had the opposite sensitization order, first receiving OVA-IMDQ+SDI twice, thus establishing a balanced Type 1/Type 2 response to OVA and then attempting to cause an allergic response by sensitizing twice with OVA-Alum (indicated hereafter as IMDQ+SDI➔Alum). To further favor the allergic phenotype, mice were challenged three times intranasally with 20 μg of OVA over a period of five days (one day on/one day off). Additionally, to investigate the role of mouse genetic background in the ability of IMDQ+SDI to revert or prevent allergic sensitization, we did the same experiment in BALB/c mice, traditionally regarded as a Type 2-skewed model, in contrast to C57BL/6 which is considered Type 1-skewed (Fig. 2A).

**Fig 2.**
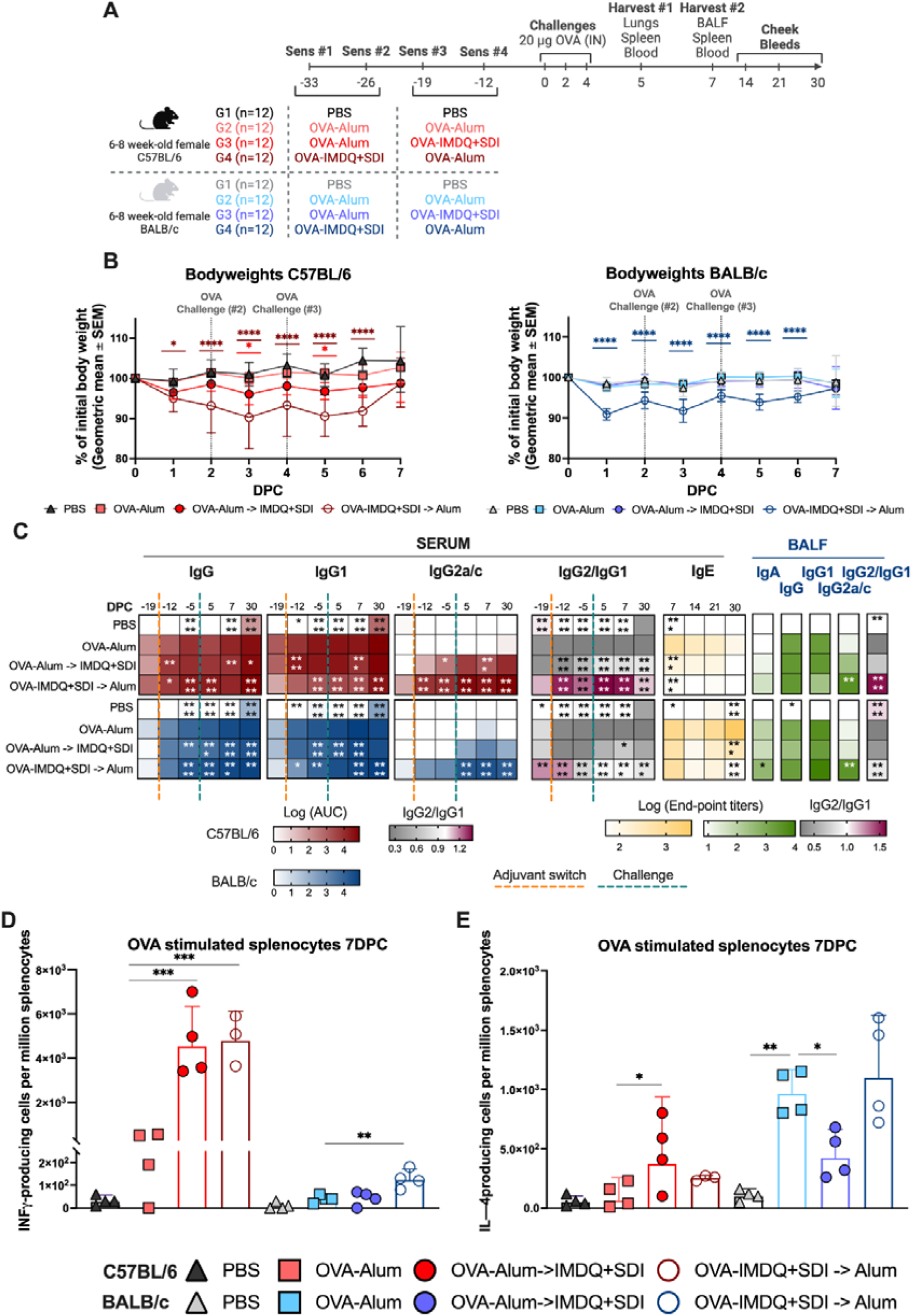
IMDQ+SDI sensitization reverts allergic antibody and T-cell responses towards a balanced Type 1/Type 2 response in OVA-Alum sensitized C57BL/6 and ameliorates them in sensitized BALB/c. A) Experimental design. Created with: <u>Biorender.com</u>. B) Daily mice bodyweights from 0DPC to 7DPC. C) Heatmaps representing antibody reponses, described form left to right: OVA-binding IgG, OVA-binding IgG1 and OVA-binding IgG2c represented as mean area under the curve (AUC) of the optical density at 450 nm (OD450) minus the background noise (OD650); C57BL6 in red and BALB/c in blue. Skewing of the antibody response represented as the coefficient between the log_10_ of the OVA-binding IgG2 (AUC) and the log_10_ of the OVA-binding IgG1 (AUC). Minimum values are set to 1 for calculation purposes. OVA-binding End-point titers of Total IgE measured in serum after OVA challenge represented as ELISA endpoint-titers. Mucosal antibody response measured in BALF at 7DPC represented as ELISA end-point titers. Statistical analysis: Kruskal-Wallis one-way ANOVA with Dunn’s multiple comparisons test using the OVA-Alum (IP) group as control for multiple comparisons. D) IFN-γ producing cells per million splenocytes measured by ELISpot at 7DPC. E) IL-4 producing cells per million splenocytes measured by ELISpot at 7DPC. Statistical analysis: for comparisons among groups at several time points: Two-way ANOVA with Tukey’s multiple comparisons test using the OVA-Alum (IP) group as control for multiple comparisons. Color of the significance markers indicates the comparison they represent; for comparisons among groups at a specific time point: Kruskal-Wallis one-way ANOVA with Dunn’s multiple comparisons test using the OVA-Alum (IP) group as control for multiple comparisons. Significance is represented as:*p = 0.05 to 0.01, **p = 0.01 to 0.001, ***p = 0.001 to 0.0001, ***p < 0.0001.

Consistent with the previous experiment, C57BL/6 mice lost significantly more weight the day after the challenge (1DPC, 3DPC and 5DPC, in this experiment) if they were sensitized with IMDQ+SDI. The IMDQ+SDI➔Alum group lost significantly more weight than the Alum group and the weight remained significantly lower from 1DPC to 6DPC. Additionally, 3 of 12 animals in that group reached humane end point before the end of the experiment. The Alum➔IMDQ+SDI group also showed significantly more weight loss after the second and third challenges with OVA (3 and 5 DPC), but animals did not reach humane endpoint. No morbidity or mortality was detected in the PBS or Alum groups. Like C57BL/6, morbidity was also detected in BALB/c after the first challenge, leading to a significant bodyweight loss of the IMDQ+SDI➔Alum group, which remained significant from 1DPC to 6DPC. Unlike C57BL/6, BALB/c recovered most of the weight lost the second day after each challenge and no mortality was detected. Additionally, the Alum➔IMDQ+SDI group did not lose significantly more weight than the Alum group (Fig. 2B).

One week after the second sensitization, regardless of the adjuvant used, all C57BL/6 mice (except the PBS group) presented detectable OVA-binding IgG titers. No significant differences were found among groups at this point. The adjuvant switch resulted in a significant antibody boost after one week in the Alum➔IMDQ+SDI group. The IMDQ+SDI➔Alum group also showed significantly higher IgG titers than the Alum group. After the fourth sensitization, only the IMDQ+SDI➔Alum group maintained significantly higher titers than the Alum group, which remained significantly higher after the 3 challenges (5DPC) and until the end of the experiment (30DPC). The Alum➔IMDQ+SDI group presented significantly higher titers than the Alum group at 7DPC, which remained higher until the end of the experiment. OVA-binding IgG titers followed a very similar profile in BALB/c (Fig. 2C).

Focusing on IgG subclasses, OVA-binding IgG1 titers were similar in the Alum and Alum➔IMDQ+SDI C57BL/6 groups after the four sensitizations and remained that way until the end of the experiment. Interestingly, the adjuvant switch from Alum to IMDQ+SDI boosted OVA-binding IgG1, leading to significantly higher titers one week after the switch. The IMDQ+SDI➔Alum group presented significantly lower OVA-binding IgG1 than the Alum control, which remained lower even after receiving two Alum doses and three OVA challenges. The PBS group that also received the three OVA challenges, showed some OVA-binding IgG1 titers at 30DPC, suggesting that even in the absence of adjuvant-favored sensitization, there is some antibody response to OVA and is skewed towards a Type 2 response. OVA-binding IgG2c titers were undetectable or extremely low in the PBS and Alum C57BL/6 groups throughout the experiment, as expected. The IMDQ+SDI➔Alum group rapidly generated detectable OVA-binding IgG2c titers, which remained significantly higher compared to the Alum control throughout the experiment. The Alum➔IMDQ+SDI group showed detectable OVA-binding IgG2c titers as soon as one week after the first dose of IMDQ+SDI. Although the titers increased after a second dose of IMDQ+SDI and the challenge, they never caught up to the level of the IMDQ+SDI➔Alum group. In all, these results show that there was some Type 1/Type 2 flexibility but priming towards a specific response had an impact in the final antibody subclass skewing. While it is possible to skew the antibody response from previously sensitized mice towards a balanced Type 1/Type 2 response, mice receiving IMDQ+SDI first tend to show a more Type 1-skewed response. Still, these differences are mitigated by 30DPC, thus order of adjuvant administration might be less important at later timepoints. Relevant differences in OVA-binding IgG subclasses were found in BALB/c compared to C57BL/6. OVA-binding IgG1 titers in the IMDQ+SDI➔Alum group rapidly reached similar levels to the Alum and Alum➔IMDQ+SDI groups in BALB/c after the adjuvant switch, while in C57BL/6 remained noticeably lower until the end of the experiment. Similar to C57BL/6, PBS controls also produce OVA-binding IgG1 at 30DPC. On the other hand, OVA-binding IgG2a titers were only detectable in the IMDQ+SDI➔Alum group prior to challenge. While the Alum➔IMDQ+SDI group eventually produces IgG2a at 5DPC, they remained noticeably lower and start to decrease by 30DPC. Overall, Alum group showed a very strong Type 2 skewing, Alum-IMDQ+SDI presented a similar profile prior to challenge, and it trended towards a more balanced antibody response, still was Type 2 skewed, at 30DPC. IMDQ+SDI➔Alum maintained a balanced Type 1/Type 2 response throughout the experiment, suggesting a reduced flexibility of the skewing in BALB/c compared to C57BL/6 and a more permanent effect of priming in the final response (Fig. 2C).

Total IgE levels in serum were the highest in both mouse strains in the Alum group. In C57BL/6, while they were reduced, IgE was still detectable in all the mice from the Alum➔IMDQ+SDI, but only in one receiving IMDQ+SDI first, suggesting again an important role of adjuvant priming in antibody production. In BALB/c, all the mice in Alum➔IMDQ+SDI and IMDQ+SDI➔Alum groups presented detectable IgE at 7DPC. Interestingly, IgE decreased in the groups receiving IMDQ+SDI, being detectable in only 2 mice from the Alum➔IMDQ+SDI group and one in the IMDQ+SDI➔Alum group by 30 DPC, while titers remained high in the Alum group (Fig 2C).

Mucosal antibody response in BALF at 7DPC shows a similar trend to the serum responses in both mouse strains with OVA-binding IgG1 higher in the Alum and Alum➔IMDQ+SDI groups and OVA-binding IgG2c or IgG2a present mostly in groups receiving IMDQ-SDI, with higher titers in the IMDQ+SDI➔Alum group. In C57BL/6, OVA-binding IgA titers could only be detected in half of mice from the Alum➔IMDQ+SDI group, while it was detectable in all animals from the IMDQ+SDI➔Alum group. BALB/c presented a stronger induction of OVA-binding IgA titers than C57BL/6, not only in the IMDQ+SDI➔Alum group, but also the Alum and Alum➔IMDQ+SDI groups (Fig 2C).

In the previous experiment, we demonstrated an elevated T-cell recruitment to the lung of mice sensitized with IMDQ and IMDQ+SDI. To assess systemic T-cell activation due to IMDQ+SDI sensitization, we performed IFN-γ and IL-4 ELISpots in splenocytes stimulated with OVA overlapping peptides at 7DPC. As expected, overall IFN-γ T-cell responses to OVA peptides were higher in C57BL/6 while IL-4 T-cell activation was higher in BALB/c. In C57BL6, both groups sensitized with IMDQ+SDI, regardless of the order of administration, presented a very robust IFN-γ producing T-cell activation when exposed to OVA overlapping peptides. Unexpectedly, while the Alum group presented some IL-4 splenocyte activation, it was significantly lower than in the Alum➔IMDQ+SDI group. Even the group IMDQ+SDI➔Alum presented higher (not significant) activation of IL-4 producing T-cells. In BALB/c, only the IMDQ+SDI➔Alum group presented significant IFN-γ producing cells. In contrast to C57BL/6, IL-4 producing cells were significantly higher in the Alum group than the Alum➔IMDQ+SDI group in BALB/s, while the IMDQ+SDI➔Alum group showed similar levels to the Alum control (Fig. 2D and E).

Cytokine profiling in the lungs of both mouse strains at 5DPC once again shows a Type 2 cytokine profile in the Alum group. These animals showed elevated IL-4, 5, 13, 9, 33 and 6 as well as GM-CSF, measured as Z-score of the average MFI in the Luminex assay, compared to the groups receiving IMDQ+SDI. In C57BL/6, IL-4 and IL-5 are significantly reduced by the administration of IMDQ+SDI, regardless of the timing. GM-CSF and IL-6 are also significantly reduced in the IMDQ+SDI➔Alum group, while the difference is not significant in the Alum➔IMDQ+SDI group. The remaining differences are not statistically significant. On the other hand, other cytokines significantly increased with IMDQ+SDI administration. In the Alum➔IMDQ+SDI group, only RANTES is significantly increased compared to the Alum group; while RANTES, IP-10, TNF-α, MIP-2α and MIP-1β were significantly elevated in the group receiving IMDQ+SDI first. Cytokine profile in BALB/c also reveals a significant reduction of IL-4 and GM-CSF in both groups receiving IMDQ+SDI compared to the Alum control, but only a non-significant reduction of IL-5 and IL-13. (Fig. 3A, Fig. S3 and S4). In contrast, only the IMDQ+SDI➔Alum group showed significantly elevated levels of Type 1 cytokines such as IL-1β, INF-γ and MIP-1β, but no significant differences in any of them were detected between the Alum and Alum➔IMDQ+SDO group. Cytokine/chemokine profiling in the spleen also suggest a broader Type 1 activation in C57BL/6 groups receiving IMDQ+SDI, than in BALB/c (Fig S5), although a certain compartmentalization of cytokine response can be found in both strains comparing lung and spleen cytokines.

**Fig 3.**
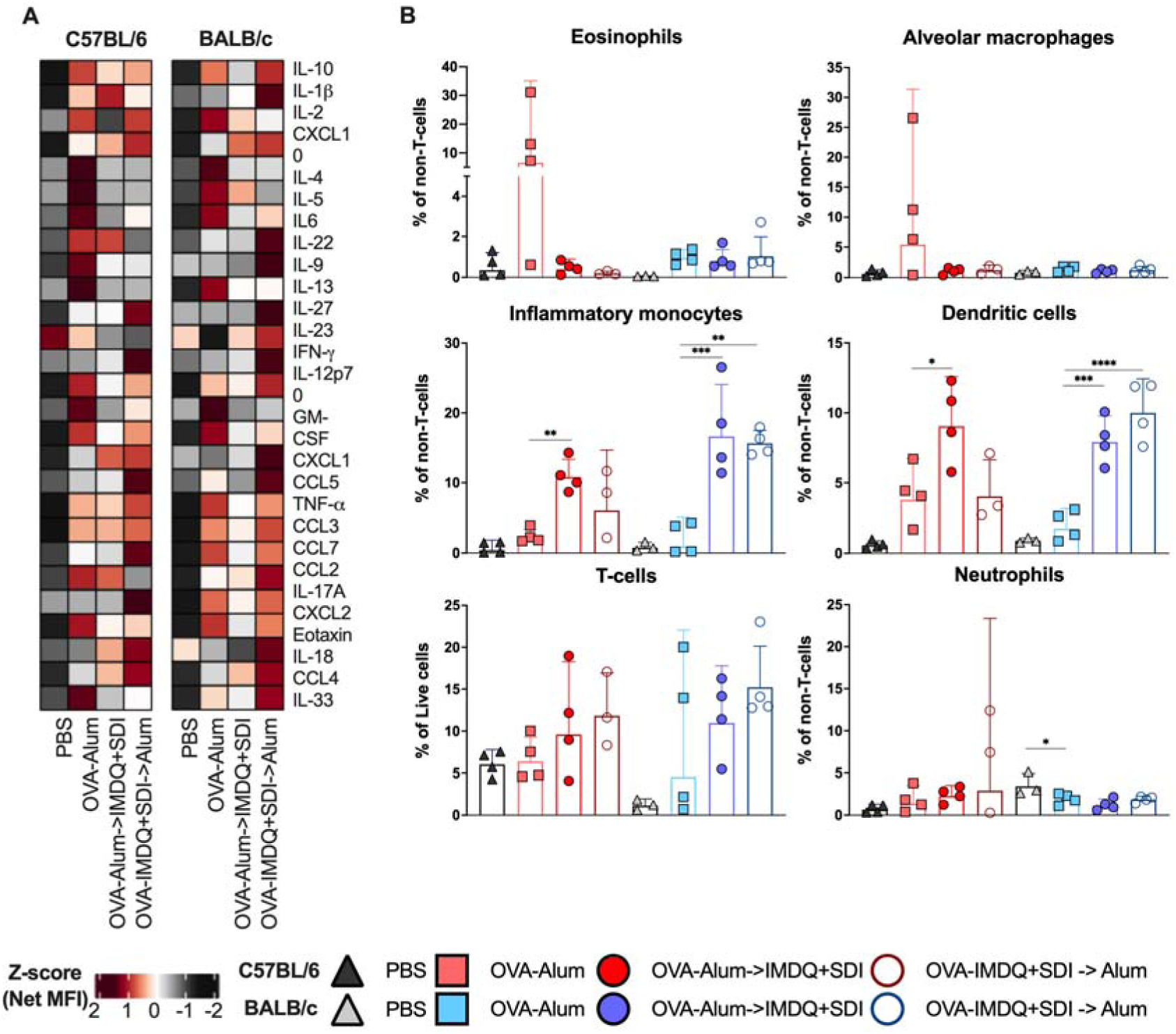
IMDQ+SDI sensitization aborts lung eosinophilia and Type 2 cytokine production after OVA challenge in C57BL/6, but does not prevent allergic phenotype in sensitized BALB/c. A) Heatmap of net mean fluorescence intensity (MFI) for 27 cytokines/chemokines, z-scored by row, measured in lung homogenates at 5DPC. B) Percentage of lung cell populations measured by flow cytometry at 5DPC. From left to right and top to bottom: Eosinophils (Gating: Live > CD3^-^ > Ly6G^-^ > CD11c^-^ CD11b^+^ > SiglecF^+^), alveolar macrophages (Gating: Live > CD3^-^ > Ly6G^-^ > CD11c^+^ CD11b^-^ > SiglecF^+^), inflammatory monocytes (Gating: Live>CD3^-^> Ly6G^-^ > CD11c^-^ CD11b^-^ > Ly6C^+^), dendritic cells (Gating: Live > CD3^-^ > Ly6G^-^ > CD11c^+^>MHCII^+^ SiglecF^-^), T-cells (Gating: Live>CD3^+^) and neutrophils (Gating: Live > CD3^-^ > Ly6G^+^ CD11b^+^). Statistical analysis: Kruskal-Wallis one-way ANOVA with Dunn’s multiple comparisons test using the OVA-Alum (IP) group as control for multiple comparisons. Significance is represented as: *p = 0.05 to 0.01, **p = 0.01 to 0.001, ***p = 0.001 to 0.0001, ***p < 0.0001.

Flow cytometry analysis of lung cell populations at 5DPC reflected important differences between mouse strains. In C57BL/6, a reduction in eosinophil recruitment was observed in those groups receiving IMDQ+SDI, regardless of the timing, as well as a reduced fraction of alveolar macrophages. Groups receiving IMDQ+SDI presented a higher recruitment of inflammatory monocytes and dendritic cells, as well as a non-significant increase of T-cells. In BALB/c similar lung eosinophilic recruitment was detected in the Alum, Alum➔IMDQ+SDI and IMDQ+SDI➔Alum groups (Fig 3B, Fig. S5). Overall, the results in BALB/c suggest a limited flexibility from Type 2 to Type 1 once mice are primed, although IMDQ+SDI leads to some reduction of IgE antibody at late time-points and therefore, might have an effect on subsequent intranasal challenges with the allergen or other antigens.

### IMDQ+SDI improves immune responses against influenza challenge in OVA-sensitized mice, regardless of their genetic background

The role of asthmatic allergy in the outcome of influenza infections is still poorly characterized^23^. Studies in mice show both aggravated morbidity of influenza infection^24^ as well as increased resistance^25^ in OVA-sensitized mice. Adjuvants are known to induce durable reprogramming of the innate immune system which could provide broad protection against diverse pathogens^26^. Thus, we hypothesized that OVA sensitization with IMDQ-SDI might prime the lung towards a faster and more efficient response against unrelated antigens, such as an influenza virus.

To test this hypothesis, surviving BALB/c and C57BL/6 mice at 30DPC from the previous experiment were challenged with a sublethal dose of H1N1 [0.2 LD_50_ of A/New Caledonia/20/1999 (NC99)]. Mice body weights were recorded daily to evaluate morbidity and at 5 days-post-infection (DPI) animals were euthanized and lungs and blood were harvested.

Both strains presented significant morbidity, measured as body weight loss, in mice from the Alum groups (mean of the percentage of initial body weight in BALB/c was 84.2% and 88.5% in C57BL/6 at 5DPI). In BALB/c, the PBS group lost the least amount of weight (96.5%) while Alum➔IMDQ+SDI and IMDQ+SDI➔Alum groups presented 92.2% and 91.2% of the initial body weight, respectively. Contrarily, C57BL/6 sensitized with IMDQ+SDI presented reduced morbidity (Alum➔IMDQ+SDI = 94.4%; IMDQ+SDI➔Alum = 95.5%) compared to the PBS group (91.6%) (Fig. 4A). Lung viral titers at 5DPI were also elevated in the Alum groups compared with the other groups for both mouse strains. In the case of BALB/c, the Alum group presented a significant 25-fold increase compared with the PBS group, while a smaller 5.3-fold increase compared to the Alum➔IMDQ+SDI group, and a 3.6-fold increase compared to the IMDQ+SDI➔Alum group. On the other hand, differences for C57BL/6 were not as dramatic for the PBS group (3.7-fold increase) and were higher in the groups receiving IMDQ+SDI (5.8-fold for Alum➔IMDQ+SDI and 4.5-fold for IMDQ+SDI➔Alum) (Fig 4B), correlating strongly with bodyweight data in both strains.

**Fig 4.**
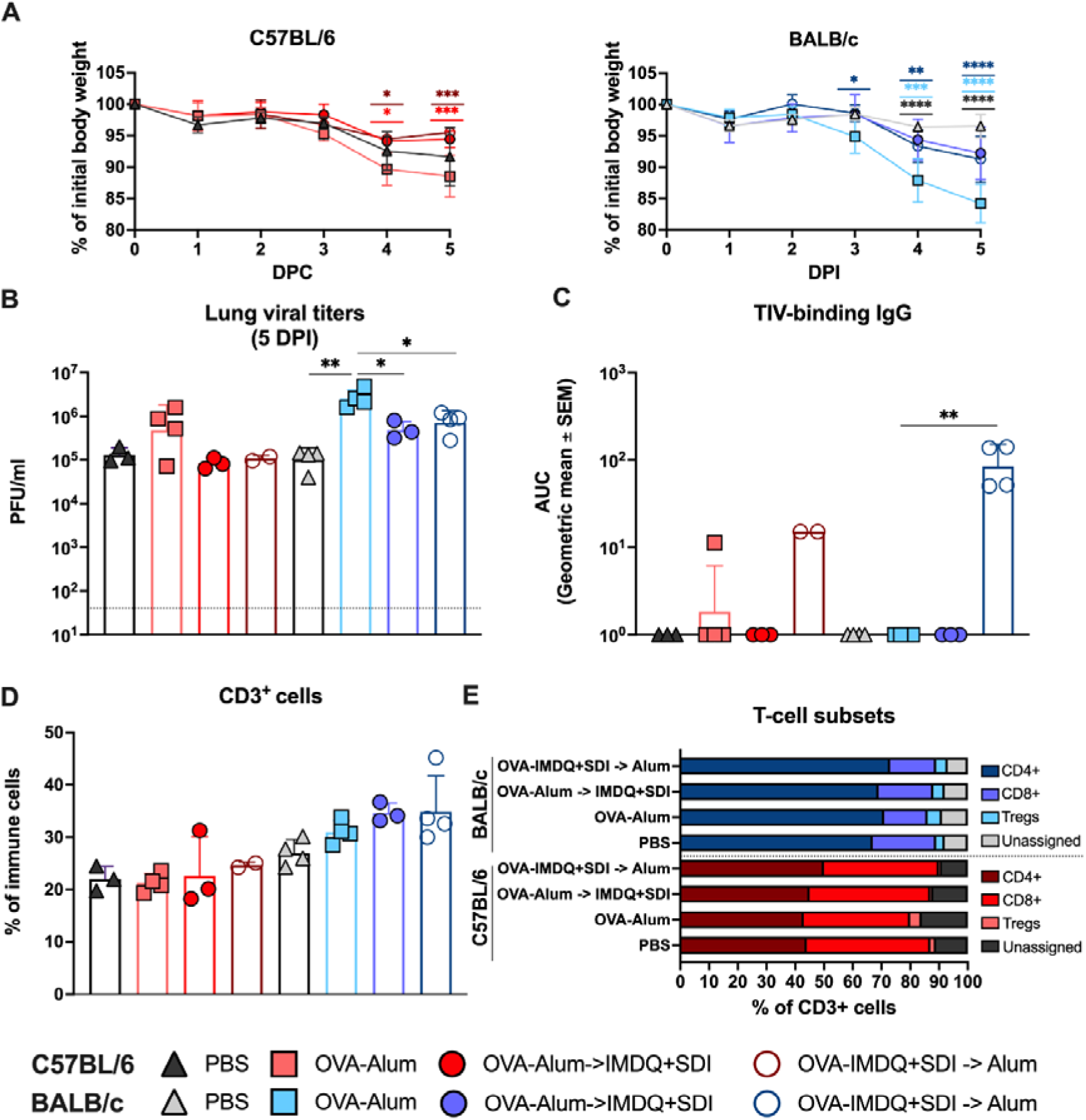
Priming or boosting both mouse strains with OVA-IMDQ+SDI leads to improved outcomes of a sublethal influenza infection. A) Daily mice bodyweights measured from 0DPI to 5DPI. B) Lung viral titers expressed as plaque forming units per mL (PFU/mL) measured in lung homogenates at 5DPI. Limit of detection = 40 PFU/mL. C) Trivalent Influenza Vaccine (containing NC99 HA)-binding total IgG measured at 5DPI represented as Area under the curve (AUC) of the optical density at 450 nm (OD450) minus the background noise (OD650). Lung cell subpopulations at 5DPI are presented in panels D to H. D) Percentage of lung total T-cells (CD3+, gating: Live> CD45^+^> CD3^+^). E) Percentage of T-cell subpopulations represented as the mean % of CD3% cells per group: CD4^+^ cells (Gating: Live> CD45^+^> CD3^+^>CD4^+^CD8^-^), CD8^+^ cells (Gating: Live> CD45^+^> CD3^+^>CD4^-^CD8^+^), T_regs_ (Gating: Live> CD45^+^> CD3^+^>CD4^+^CD8^-^>FoxP3^+^). CD3^+^ cells not included in any of the three subpopulations are categorized as “unassigned”. Statistical analysis: for comparisons among groups at several time points: Two-way ANOVA with Tukey’s multiple comparisons test using the OVA-Alum (IP) group as control for multiple comparisons. Color of the significance markers indicates the comparison they represent; for comparisons among groups at a specific time point: Kruskal-Wallis one-way ANOVA with Dunn’s multiple comparisons test using the OVA-Alum (IP) group as control for multiple comparisons. Significance is represented as:*p = 0.05 to 0.01, **p = 0.01 to 0.001, ***p = 0.001 to 0.0001, ***p < 0.0001.

Although 5DPI is too early to detect meaningful NC99-binding antibodies, we performed ELISAs to measure Trivalent Influenza Vaccine (TIV)-binding IgG (Vaccine containing NC99 HA). Interestingly, although IgG titers were very low, they were already detectable at 5DPI in the IMDQ+SDI➔Alum groups, while they were not detectable in any of the other groups (Fig 4C). Antibody response against OVA was not greatly affected by the H1N1 challenge, which is relevant as the virus is produced in eggs. Trends observed at 30 DPC continue after NC99 challenge at 5DPI, with similar titers of each antibody subtype. In BALB/c, OVA-binding IgG2a continued to wane down in the Alum➔IMDQ+SDI group, while this was not the case for C57BL/6. Again, BALB/c Alum group showed significant IgE, that could also be detected in the groups receiving IMDQ+SDI to a lesser extent, while C57BL/6 presented IgE only in the group sensitized with Alum OVA only (Fig. S7).

Then we focused on the analysis of T-cell, macrophage, neutrophil and eosinophil subpopulations in the lung to evaluate the effect of the sensitization in the cellular response against NC99. The overall number of T-cells showed no significant differences among groups for BALB/c, while increased numbers were found in the Alum and Alum➔IMDQ+SDI groups of C57BL/6, although these increases did not lead to elevated percentages of immune cells. For the Alum group, most of the T-cell increase was driven by elevated numbers of T_regs_, while in the case of the IMDQ-Alum group both CD8^+^ and CD4^+^ subsets were significantly elevated. In all cases, these differences were only statistically significant in terms of total number of cells, but not percentage of T-cells. In BALB/c, both PBS and Alum➔IMDQ+SDI present increased numbers of CD8^+^ T-cells compared to the Alum group, T_regs_ were elevated in all groups compared to the PBS, while no differences were found among groups in total CD4^+^ T-cell count. Contrary to C57BL/6, CD4^+^ T-cell showed significant differences when evaluated as a fraction of CD3^+^, with the Alum and IMDQ+SDI➔Alum groups showing significantly higher proportions of CD4^+^ (Fig 5D and E, S8).

**Fig 5.**
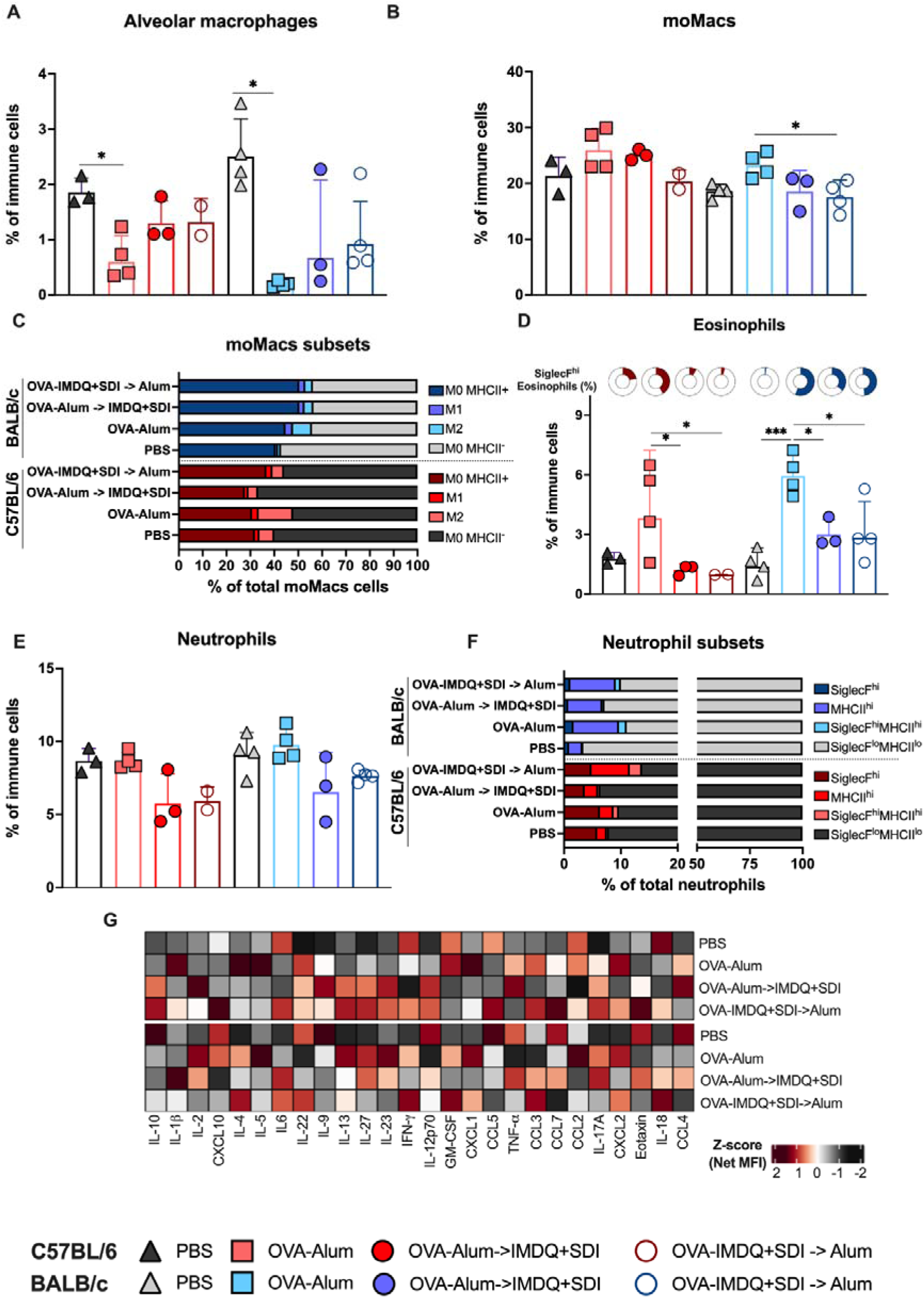
Priming or boosting mice with OVA-IMDQ+SDI led to a reduction in SiglecF expression in lung eosinophils and neutrophils and the reduction of IL-5 after NC99 challenge compared to allergic mice. A) Percentage of immune cells identified as alveolar macrophages (Gating: Live> CD45^+^> Ly6G^-^> SiglecF^+^CD11c^+^). B) Percentage of total monocyte-derived macrophages (moMacs) in the lung at 5DPC (Gating: Live> CD45^+^> Ly6G^-^> SiglecF^-^>Ly6C^+^F4/80^+^). C) Distribution of moMacs subpopulations: M0 MHCII^+^ (Gating: Live> CD45^+^> Ly6G^-^> SiglecF^-^>Ly6C^+^F4/80^+^> MHCII^+^> ArgI^-^iNOS^-^), M1 moMacs (Gating: Live> CD45^+^> Ly6G^-^> SiglecF^-^>Ly6C^+^F4/80^+^> MHCII^+^> ArgI^-^iNOS^+^), M2 moMacs (Gating: Live> CD45^+^> Ly6G^-^> SiglecF^-^>Ly6C^+^F4/80^+^> MHCII^+^> ArgI^+^iNOS^-^) and M0 MHCII^-^ (Gating: Live> CD45^+^> Ly6G^-^> SiglecF^-^>Ly6C^+^F4/80^+^> MHCII^-^). D) Percentage of eosinophils (in barplot) and fraction of those eosinophils in each group classified as SiglecF^hi^ (donut charts). Total Eosinophils (Gating: Live> CD45^+^> Ly6G^-^> SiglecF^+^CD11b^+^) and SiglecF^hi^ eosinophils (Gating: Live> CD45^+^> Ly6G^-^> SiglecF^hi^CD11b^+^). E) Percentage of total neutrophils: (Gating: Live> CD45^+^> Ly6G^+^> SiglecF^+^CD11b^+^). F) Distribution of neutrophil subpopulations. SiglecF^hi^ neutrophils (Gating: Live> CD45^+^> Ly6G^+^> SiglecF^+^CD11b^+^>SiglecF^+^) and MHCII^hi^ neutrophils (Gating: Live> CD45^+^> Ly6G^+^> SiglecF^+^CD11b^+^> MHCII^+^), SiglecF^hi^MHCII^hi^ neutrophils (Gating: Live> CD45^+^> Ly6G^+^> SiglecF^+^CD11b^+^>SiglecF^+^MHCII^+^) and SiglecF^lo^MHCII^lo^ neutrophils (Gating: Live> CD45^+^> Ly6G^+^> SiglecF^+^CD11b^+^>SiglecF^-^MHCII^-^). G) Heatmap of net mean fluorescence intensity (MFI) for 26 cytokines/chemokines, z-scored by column, measured in lung homogenates at 5DPI. Statistical analysis: Kruskal-Wallis one-way ANOVA with Dunn’s multiple comparisons test using the OVA-Alum (IP) group as control for multiple comparisons. Significance is represented as:*p = 0.05 to 0.01, **p = 0.01 to 0.001, ***p = 0.001 to 0.0001, ***p < 0.0001.

H1N1 infection normally leads to a significant depletion of alveolar macrophages in the lung that peaks between 5 and 7 DPI. In the Alum group a significantly lower percentage of alveolar macrophages was detected, while in groups receiving IMDQ+SDI were partially protected from this loss (Fig 5A). Four monocyte-derived macrophages (moMacs) subsets were defined based on their expression of MHC-II only (M0 MHCII^+^), iNOS (M1 or pro-inflammatory) or Arg-I (M2 or anti-inflammatory). Cells Ly6C^+^F4/80^+^MHCII^-^ cells were defined as M0 MHCII^-^. Overall, the Alum group presented elevated fractions of moMacs, that correlates with the alveolar macrophage depletion. This increase in moMacs was significant when compared to IMDQ+SDI➔Alum BALB/c mice (Fig. 5B). For both strains, the majority of moMacs were defined as M0, and no significant differences were found among groups in the fraction of moMacs they represented. Interestingly, in BALB/c M1 moMacs comprised a significantly higher fraction in the Alum and IMDQ+SDI➔Alum groups compared to the PBS group. A similar trend was found for C57BL/6 but differences were not significant. M2 macrophages were significantly elevated in the Alum group compared to the other three for BALB/c. Once again, while a similar trend could be appreciated in C57BL/6, differences were not significant (Fig 5C, S8).

Recently, activation of eosinophils has been associated with higher expression of Siglec-F (Siglec-F^hi^ eosinophils) and they may have an important role, especially in the context of breakthrough viral infections^27,28^. It is also suggested that eosinophils can act as APCs in upper respiratory airways allergic reactions^29^. Therefore, we also investigated lung eosinophil subpopulations based on their Siglec-F and MHC-II expression. Total number of eosinophils in the lung was elevated in both mouse strains in the Alum group. In C57BL/6, Siglec-F^hi^ eosinophils were significantly elevated in the Alum group, while the groups that received IMDQ+SDI presented levels lower than the PBS group. On the other hand, while the Siglec-F^hi^ eosinophils are significantly reduced in the Alum➔IMDQ+SDI and IMDQ+SDI➔Alum groups for BALB/c compared to the Alum group, they were elevated compared to the PBS group (Fig 5D). No significant differences were found in either of the mouse strains in MHC-II^+^ eosinophils.

The neutrophil fraction in the lung trended towards higher in both mouse strains in the PBS and Alum groups, although not significantly (Fig. 5E). Similar to eosinophils, a subpopulation of Siglec-F^+^ Neutrophils, which presents an activated phenotype, has been shown to increase in an allergic rhinitis model^30^. Additionally, some MHC-II and co-stimulatory molecules have been shown to be induced on neutrophils, which may serve as antigen presenting cells (APC) for T-cells in an allergic context^31^. Thus, we investigated these neutrophil subsets in the context of a viral infection in previously OVA-sensitized mice. In both strains, the fraction of Siglec-F^+^ neutrophils was significantly higher in the Alum group. On the other hand, MHC-II^+^ neutrophils were only elevated in the IMDQ+SDI➔Alum group for C57BL/6 while all groups present an elevated fraction for BALB/c when compared to the PBS group (Fig 6F).

**Fig 6.**
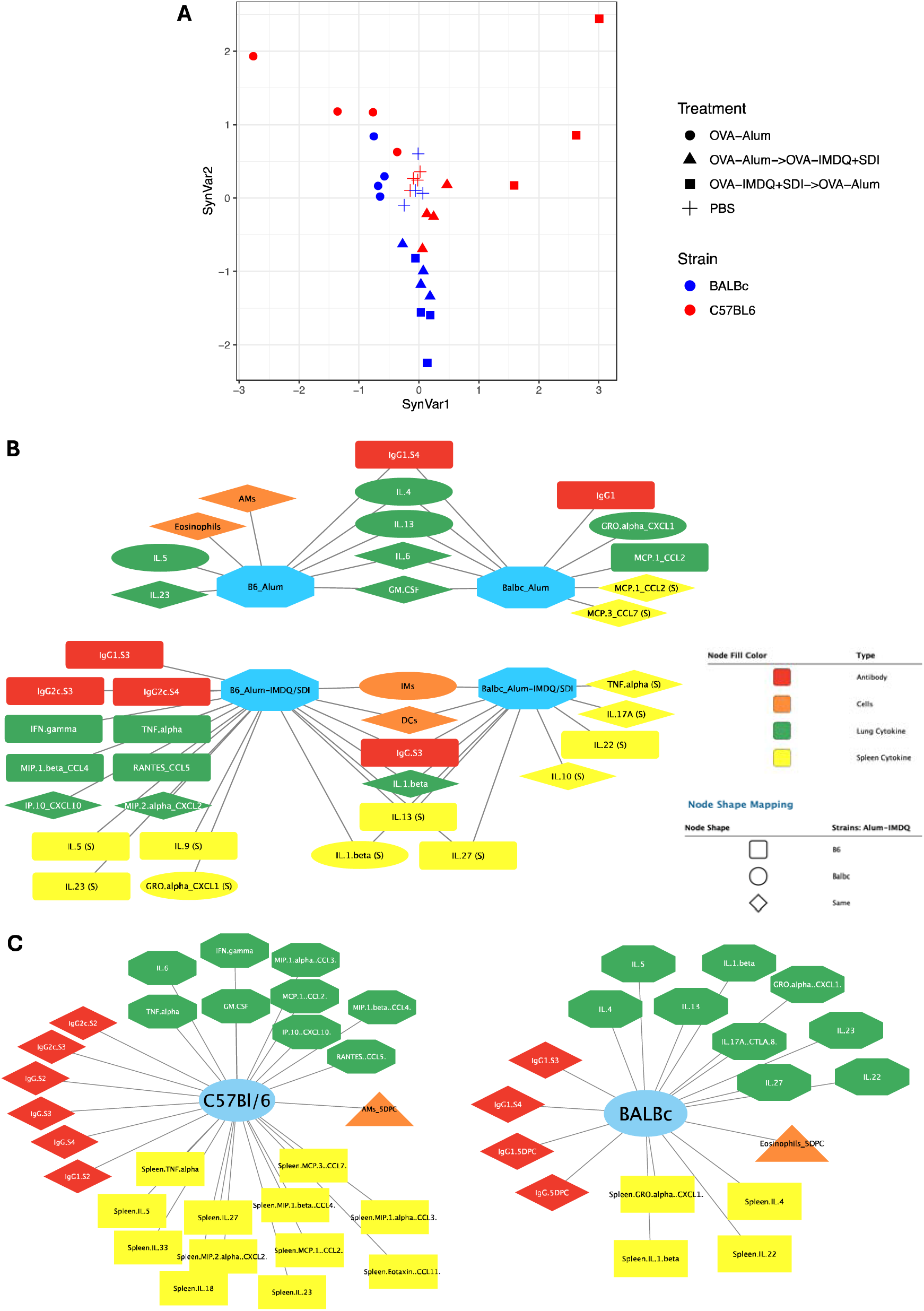
Multiple co-inertia analysis reveals divergent effects of IMDQ+SDI in OVA-sensitization in BALB/c versus C57BL/6 5 days after OVA challenge. Multiple co-inertia analysis (MCIA) was performed for multi-Omic integration and network modeling to identify the antibody, cytokine and cellular features that distinguish adjuvant treatments within and across mouse strains. A) PCA plot of SynVar1 and SynVar2, the two largest drivers of variance in the multi-Omic dataset, showing separation of samples by strain and by adjuvant skewing between and within strains. B-C) Cytoscape was used to generate network models identifying the top features from SynVar1 and SynVar2 associated with each distinct adjuvant treatment and mouse strain. To generate networks, we identified the components of SynVar1 and SynVar2 which were significantly associated to B) Alum vs Alum-IMDQ for both strains and C) Alum-IMDQ comparing C57Bl/6 and BALB/c. Features are linked to the adjuvant conditions and strain they are significantly associated to, identifying critical differences in the types of immune resposnes generated across mouse strains for these different adjuvant skewing conditions.

Lung cytokine/chemokine profiling at 5DPI revealed interesting differences among groups and between mouse strains (Fig 5G). However, most differences were small and non-significant in terms of protein concentration. Only a significant reduction in IL-22 could be detected in Alum➔IMDQ+SDI C57BL/6 mice when compared to the Alum group (Fig. S9 and S10). Still some relevant trends could be highlighted. In the Alum groups, NC99 infection led to the increased production of IL-4 and IL-5. These cytokines were at the level of the PBS group for any of the groups receiving IMDQ+SDI in C57BL/6 mice, but only IL-5 was reduced in those groups of BALB/c. IL-9 and GRO-α had a similar profile to IL-4, but only on C57BL/6. Interestingly, some cytokines were elevated in both the Alum group and the IMDQ+SDI➔Alum, but not in the Alum➔IMDQ+SDI, including IL-23, IFN-γ, CCL-3, CCL-7, CCL-2, MIP-2α or IL-18.

### Multiple co-inertia analysis reveals divergent effects of IMDQ+SDI in OVA-sensitization in BALB/c versus C57BL/6, but a similar effect on the outcome of NC99 challenge

To further establish correlates between antibody, cytokine/chemokine profiles and cellular recruitment and activation; and to evaluate differential responses based on mouse genetic background, we performed a multiple co-inertia analysis (MCIA)^32^. This method is an exploratory data analysis method that allowed us to identify co-relationships between multiple high dimensional datasets (antibody response, cytokines/chemokines and cell frequencies), projecting them into the same dimensional space and transforming them onto the same scale. After this processing, the method allowed for the identification of the most relevant variants from each dataset. MCIA of mice 5 days after OVA challenge (5DPC) (Fig 6A) revealed that most variance in the data could be explained by differences among the different C57BL/6 groups, specially the IMDQ+SDI➔Alum group and the Alum group (SynVar1), with samples clustering together based on the four experimental groups. BALB/c mice also clustered together based on experimental group, although differences were smaller. The other main driver of variance in the data set was genetic background (SynVar2). Then we explored the specific variables that drove those differences. When Alum alone groups, were compared to Alum➔IMDQ+SDI, we could identify common drivers in both mice strains. Alum group for both BALB/c and C57BL/6 presented elevated levels of OVA-binding-IgG1 after 4 sensitizations, as well as lung cytokines IL-4, 6, 13 and GM-CSF. On the other hand, Alum➔IMDQ+SDI presented significantly higher of total OVA-binding IgG after the third sensitization, revealing a rapid effect of the adjuvant switch in both strains. In both strains, Alum➔IMDQ+SDI induced differences locally in the lung –significantly elevated fractions of inflammatory monocytes and dendritic cells as well as cytokine IL-1β– as well as systemically in the spleen –elevated IL-1β, 13 and 27–. Though we identified these communalities between mouse strains, effects of the IMDQ+SDI sensitization in the Alum➔IMDQ+SDI were much more pronounced in C57BL/6 with increased levels of OVA-binding IgG2c antibodies after the third and fourth sensitization, as well as a pro-inflammatory lung cytokine profile (IFN-γ, TNF-α, CCL4 and 5 and CXCL2 and 10). Interestingly, IL-5, 9 and 23 were associated with Alum➔IMDQ+SDI C57BL/6 in the spleen (Fig. 6B). In all, these co-relationships between Alum and Alum->IMDQ+SDI revealed a correction of allergic phenotype by IMDQ+SDI sensitization in both strains, although significantly more pronounced in C57BL/6. The largest differences between strains were found between the IMDQ+SDI➔Alum groups. C57BL/6 were characterized by high overall OVA-IgG response, and particularly IgG2c in contrast with IgG1 in BALB/c. Lung cytokine/chemokine profile in C57BL/6 revealed a strong Type 1 proinflammatory profile with elevated IFN-γ, TNF-α, CCL2-5, IL-6 or GM-CSF also associated with elevated levels of alveolar macrophages; while lung BALB/c were characterized by Type 2 cytokines such as IL-4, 5 and 13 and some associated with allergic lung pathology such as IL-1β, CXCL1 or IL-17A as well as elevated lung eosinophils (Fig 6C).

After NC99 challenge, MCIA revealed that, once again, most variance in the data could be explained by differences among the different C57BL/6 groups, specially the IMDQ+SDI➔Alum group and the Alum group (SynVar1) and genetic background (SynVar2) (Fig S11A). Alum groups of BALB/c and C57BL/6 presented a similar profile with elevated Eosinophils, SiglecF^hi^-and MHCII^hi^-neutrophils and M2 moMacs, as well as IL-5, CCL2 and CXCL2. Alum➔IMDQ+SDI groups in both mouse strains were characterized by a reduced depletion of alveolar macrophages after NC99 challenge, as well as an increased lung T-cell recruitment (CD3^+^ in BALB/c and CD8^+^ in C57BL/6) (Fig S11B). Once again, the most striking differences between strains were found in the IMDQ+SDI➔Alum groups. C57BL/6 presented were characterized by higher OVA-binding IgG2c levels and total OVA-binding IgG while BALB/c were associated with total IgE and, interestingly, TIV-binding IgG after NC99 challenge. Lung immune cell populations in C57BL/6 were represented by alveolar macrophages, M2 MoMacs, activated neutrophils (SiglecF^hi^) and MHCII^hi^ eosinophils, as well as CD8^+^ T-cells; on the other hand, BALB/c were characterized by total eosinophils and their SiglecF^hi^ subpopulation, MHCII^hi^ neutrophils and CD4^+^ T-cells (Fig S11C). Overall, MCIA analysis suggests that IMDQ+SDI has a relevant effect on OVA challenge and subsequent challenge with an unrelated antigen (NC99) in both mouse strains regardless of the timing of administration (before or after Alum). Still, the magnitude of the effect is higher in C57BL/6 than in BALB/c and is also higher if the IMDQ+SDI is administered first and then the Alum than if the Alum is administered first. Finally, while prevention of the allergic phenotype by IMDQ+SDI is more successful in C57BL/6 as highlighted by previous results, a certain degree of protection is also achieved in BALB/c, that has a long-lasting effect in subsequent unrelated antigenic challenges.

### Allergic protection conferred by IMDQ+SDI is not driven by IgG2c or IgG2b alone, CD4^+^ T-cells are needed to restrain eosinophil recruitment and CD8^+^ T-cells to inhibit eosinophil activation

The multivariate analysis performed with the data from the previous experiments suggest a role of OVA-binding IgG2c antibodies, induced by OVA-IMDQ+SDI sensitization in the protection against allergy. However, antibody subclasses IgG2a and IgG2b from BALB/c have been associated with aggravated anaphylaxis^33^. To investigate the role of IgG2b and IgG2c in the allergic process, fifteen 6–8-week-old female C57BL/6 were sensitized intramuscularly twice, one week apart, with 20 µg of OVA in combination with Alum. Three hours prior to the challenge, mice were passively transferred 200 μL of either PBS (n=5), IgG2b (n=5) or IgG2c (n=5). IgG2b and IgG2c were purified from sera collected in all prior experiments, with a Protein-A sepharose column and diluted in sterile PBS for administration (Fig S12). Then mice were challenged 3 times intranasally with 20 μg of OVA over a period of 5 days (one day on/one day off). Mice body weights were monitored daily until 5DPC and then mice were euthanized, and lungs and blood were collected (Fig S13A).

First, morbidity was assessed after the challenge. Both groups that received the antibody transfer showed increased body weight loss compared to the control after each challenge. Nevertheless, those differences were not statistically significant (Fig. S13B).

After the challenge, we focused on immune cellular responses in the lung and the activation/maturation status of both eosinophils and neutrophils, which are known key players in asthmatic allergy, and we have shown in the prior experiments are altered after IMDQ+SDI sensitization. Alveolar macrophages were depleted in groups receiving IgG2b and IgG2c, however, differences were not statistically significant (Fig. S13C). Total neutrophils and eosinophils were increased in the group receiving the transfer of IgG2b and followed a similar trend in the group receiving IgG2c, although differences were not significant (Fig. S12D and E). Neutrophils presented no significant differences among groups in phenotype markers CD101 or Siglec-F (Fig. S12F-I). Eosinophils did not reveal any significant differences in terms of Siglec-F expression (Fig. S12J and K). In all, these results suggest a limited role of IgG2c or IgG2b induced by IMDQ+SDI by themselves in the allergy outcome, while pointing towards an overall increased morbidity, eosinophilia and neutrophilia but no changes in the activation or maturation of these cell subsets.

While antibodies induced by IMDQ+SDI by themselves could be insufficient to prevent allergy, their role in combination with T-cells could provide further mechanistic insight into IMDQ+SDI induced protection from allergy. To investigate this, twenty 6–8-week-old female C57BL/6 were sensitized intramuscularly twice, one week apart, with 20 µg of OVA in combination with IMDQ+SDI. 3 and 1 days prior to the first challenge, no-depletion (n=5), CD4^+^-depletion (n=5), CD8^+^-depletion (n=5) or both CD4^+^/CD8^+^-depletion (n=5) were performed. Then mice were challenged 3 times intranasally with 20 µg of OVA over a period of 5 days (one day on/one day off). Mice body weights were monitored daily until 5DPC and then mice were euthanized, and lungs and blood were collected (Fig. 7A).

**Fig 7.**
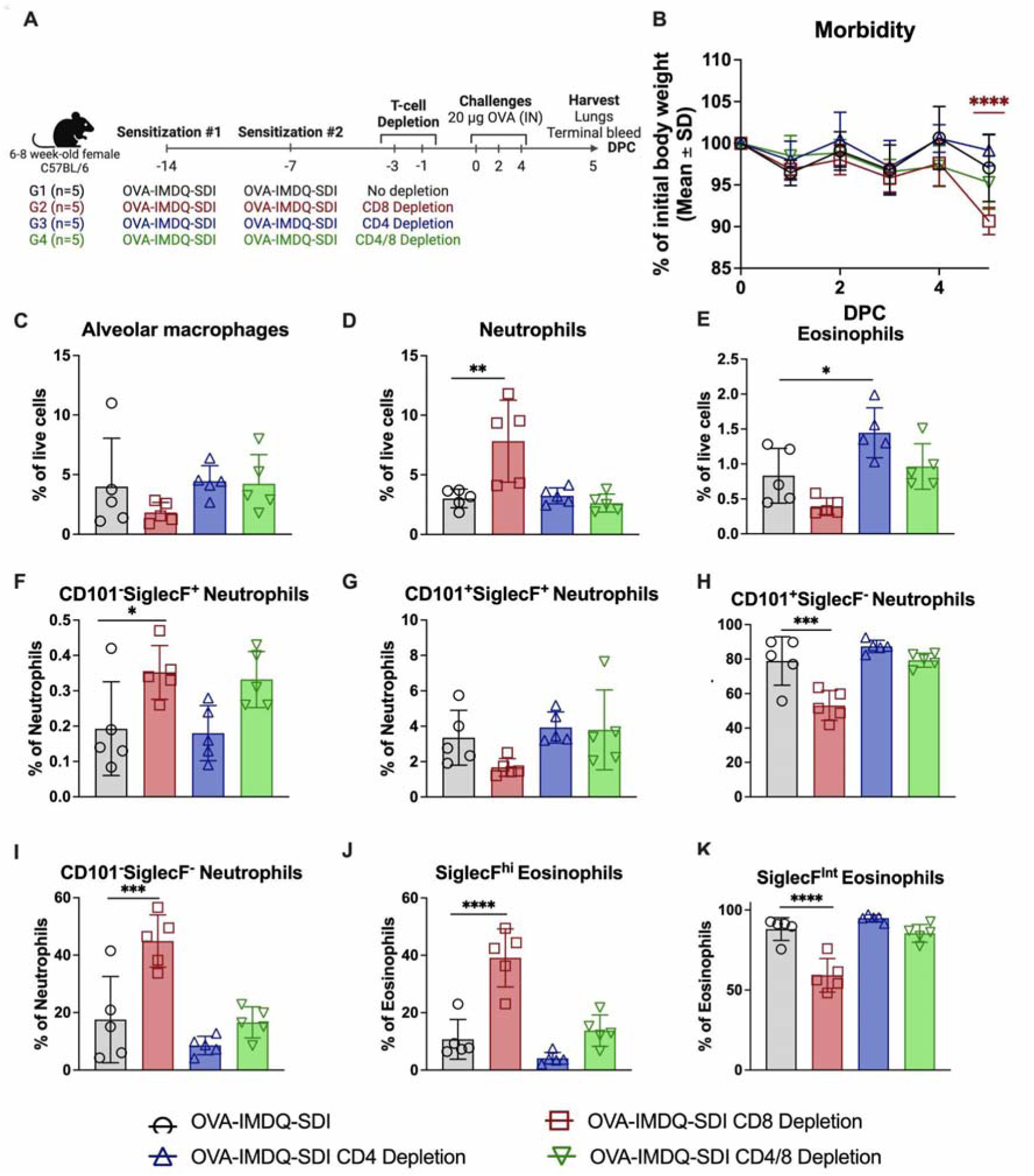
Role of CD4^+^ and CD8^+^ T-cells in the recruitment and activation of neutrophils and eosinophils in IMDQ+SDI sensitized mice. A) Experimental design. Created with: <u>Biorender.com</u>. B) Daily mice bodyweights measured from 0DPC to 5DPC. Lung flow cytometry percentage of cell subpopulations at 5DPC (C-K). C) Alveolar macrophages (Gating: Live>TER-119^-^>Ly6G^-^>CD11c^+^SiglecF^+^>MHCII^+^). D) Total neutrophils (Gating: Live>TER-119^-^>Ly6G^+^>IL-5Ra^+^CD11b^+^). E) Total eosinophils (Gating: Live>TER-119^-^ >Ly6G^-^>CD11b^+^> IL-5Ra^+^SiglecF^+^). F) CD101^-^SiglecF^+^ Neutrophils (Gating: Live> TER-119^-^>Ly6G^+^>IL-5Ra^+^CD11b^+^>CD101^-^SiglecF^+^). G) CD101^+^SiglecF^-^ Neutrophils (Gating: Live> TER-119^-^>Ly6G^+^>IL-5Ra^+^CD11b^+^>CD101^+^SiglecF^-^). H) CD101^+^SiglecF^+^ Neutrophils (Gating: Live> TER-119^-^> Ly6G^+^> IL-5Ra^+^CD11b^+^> CD101^+^SiglecF^+^). I) CD101^-^SiglecF^-^ Neutrophils (Gating: Live>TER-119^-^>Ly6G^+^>IL-5Ra^+^CD11b^+^>CD101^-^SiglecF^-^). J) SiglecF^hi^ Eosinophils (Gating: Live>TER-119^-^>Ly6G^-^>CD11b^+^> IL-5Ra^+^SiglecF^+^>CD101^+^SiglecF^hi^). K) SiglecF^int^ Eosinophils (Gating: Live>TER-119^-^>Ly6G^-^>CD11b^+^> IL-5Ra^+^SiglecF^+^>CD101^-^SiglecF^int^). Statistical analysis: for comparisons among groups at several time points: Two-way ANOVA with Tukey’s multiple comparisons test using the OVA-Alum (IP) group as control for multiple comparisons. Color of the significance markers indicates the comparison they represent; for comparisons among groups at a specific time point: Kruskal-Wallis one-way ANOVA with Dunn’s multiple comparisons test using the OVA-Alum (IP) group as control for multiple comparisons. Significance is represented as: *p = 0.05 to 0.01, **p = 0.01 to 0.001, ***p = 0.001 to 0.0001, ***p < 0.0001.

First, morbidity was assessed after the challenge. None of the groups showed significant differences after the first or second OVA challenge when compared to the non-depleted control. Interestingly, at 5DPI, mice with depleted CD-8^+^ T-cells showed a significantly higher weight loss than the non-depleted control, suggesting a potential protective role of CD8^+^ T cells against IMDQ-SDI sensitized morbidity after OVA challenge (Fig. 7B).

After the challenge, we focused on immune cellular responses in the lung and the activation/maturation status of both eosinophils and neutrophils, which we have shown in prior experiments to be altered in IMDQ+SDI groups compared to allergic groups. T cell depletion did not cause any differences in the lung alveolar macrophage populations (Fig. 7C). Total neutrophils and eosinophils had opposite outcomes depending on the depletion. CD8^+^-depletion led to an increase in neutrophils (Fig. 7D), while CD4^+^-depletion or the depletion of both T-cell subsets had no effect on the total neutrophil population (Fig 7E). The phenotype of the recruited neutrophils in CD8^+^-depleted mice corresponded to a vast majority of immature neutrophils (CD101^-^). No differences amongst groups were found in mature (CD101^+^) activated (SiglecF^+^) neutrophils (Fig. 7F-I). On the other hand, CD4^+^-depletion led to eosinophilic infiltration, while eosinophils were reduced in CD8^+^-depleted mice. Interestingly, depletion of both T-cell subsets led to an intermediate scenario. Although CD8^+^-depletion resulted in similar eosinophil levels to the PBS control, those eosinophils are highly activated (SiglecF^hi^), while almost all eosinophils in the PBS, CD4^+^-depleted and CD4^+^/CD8^+^-depleted eosinophils were SiglecF^Int^ (Fig. 7J and K). Overall, these results suggest that CD4^+^ T-cells, or a subset of them, might be responsible for restraining eosinophil recruitment in IMDQ+SDI sensitized mice while CD8^+^ T-cells, or a subset of them, limit eosinophil activation.

## DISCUSSION

Adjuvants are a key component of many vaccines, used to improve the magnitude, breadth and durability of the immune response they induce, allowing for dose sparring and improving efficacy in immunocompromised populations^34,35^. While quantitatively enhancing vaccine response is of central importance, as many purified vaccine antigens are often poorly immunogenic^36^, improving quality of the response is also a critical role of vaccine adjuvants. Specifically, quality of the response can be defined as activation of functionally optimal compartments of the immune response, such as appropriate T-cells subsets (Th1, Th2 and/or Th17) and antibody subtypes (in mice, IgG1 vs IgG2), recruitment and activation of important innate immune cell types, specific cytokine/chemokine signaling or activation of immune responses at relevant mucosal surfaces. The ability of adjuvants to change the quality of vaccine immune responses is probably most necessary in allergen immunotherapy, in which a typical strong Type 2 allergic response to an allergen is dramatically shifted, via immunization with the antigen in combination with a Type 1-skewing adjuvant, like a TLR4 or TLR9 agonist^37^.

We have shown before that a combination adjuvant comprised of a TLR7/8 agonist (IMDQ) and a RIG-I agonist (SDI) enhances influenza and SARS-CoV-2 vaccine responses in mice, yielding highly Type 1-biased vaccine immune responses^14,17^. To gain a deeper understanding on the role of this combination adjuvant in vaccine-response improvement and test it under the most stringent conditions, we set out to evaluate its immune enhancing and polarizing ability via immune profiling of pre-clinical models sensitized to the model antigen OVA, which is also used as an allergen in pre-clinical models in the context of AIT.

In mice, IgG1 is normally used a surrogate measurement of Th2 immune response, while Th1 is normally associated with increased IgG2a levels^38^. However, C57BL/6 present extremely low levels of IgG2a in serum and, normally, the antibody subtype IgG2c is selected as surrogate indicator of Th1 response^39^. Production of IgG2c is also key for the activation of other immune compartments, as it has been proved superior to other IgG subtypes in its high capacity to bind to all activating Fcγ-receptor, achieving maximal activation of monocytes, macrophages, and neutrophils^40^. We first demonstrated that naïve C57BL/6 mice sensitized with OVA-IMDQ or OVA-IMDQ+SDI produced reduced levels of OVA-binding IgG1 compared to the OVA-Alum groups, while they presented high levels of OVA-binding IgG2c antibodies, which were not detectable in the OVA-Alum groups. Several adjuvants have proved to be able to skew anti-OVA immune responses towards Type 1 or mixed Type 1/Type 2 responses in naïve mice, including TRL-9 agonist CpG ODN^41^, TLR4 agonist MPL^42^, OK-432 (a lyophilized mixture of group A *Streptococcus pyogenes*)^43^, paclitaxel^44^ or silica nanoparticles^45^. Even other TLR7/8 agonist, such as R-848, shown the ability to induce OVA-specific IgG2a in BALB/c mice, preventing allergic sensitization^46,47^. However, studies in pre-clinical models that evaluate RIG-I agonist for AIT are lacking. Other RIG-I agonists, such as 3pRNA, have shown in combination with OVA to induce cross-priming of OVA-specific CD8^+^ T-cells, using OVA as a model antigen to evaluate anticancer vaccines^48^, but was not evaluated in the context of AIT. In addition to the differential antibody subclass induction, we show that OVA-specific antibody response induced in IMDQ or IMDQ+SDI sensitized mice, was more durable than the one of Alum sensitized mice, as demonstrated by significant differences in antibody response at 11 DPC, suggesting long lasting protection against allergic reactions upon new OVA challenges. A durable response could be fundamental in AIT, as biased responses to the allergen normally wane down quickly, requiring the repeated administration of allergen extracts or products over several years^37^.

While results discussed above are relevant, the first experiment was performed in naïve mice, and it is well known that immune priming can have a dramatic effect on the ability of a given vaccine booster or adjuvant to the re-shape the immune response^49,50^. Moreover, it is unclear if Type 2-primed individuals that have received an Alum-adjuvanted vaccine can be redirected towards Type 1 by boosting with a Type 1-polarizing vaccine adjuvant, or if this would simply boost the Type 2-response the individual was primed for. In the literature, cases of both success and failure can be found. For example, Hepatitis B vaccination with a nanoemulsion adjuvant in previously Type 2-primed animals, showed increased Type 1 responses and IL-17 and decreased IL-4, IL-5 and IgG, also inducing T_regs_ and IL-10, that was required for Type 2 suppression^51^. Specifically against OVA, the oral administration of the plant compound Zerumbone, has also shown potential to skew towards Type 1 the response in previously OVA-Alum sensitized mice^52^ On the other hand, a study performed in BALB/c has shown that two heterologous boosters with Type 1-oriented CpG ODN-adjuvanted S1-based SARS-CoV-2 subunit vaccine in mice primed with two doses of Alum-adjuvanted S1-based SARS-CoV-2 subunit vaccine, were not able to revert the previously established Type 2 skewing^53^. In humans, studies in donors primed with acellular *Bordetella pertussis* vaccine (Type 2-skewing) or whole-bacteria formulation (Type 1-skewing), showed an induction of functionally different T-cell responses to the bacteria, which become fixed and remain unchanged even upon boosting^54^. To assess if Alum-primed sensitization could be reverted by boosting with a different adjuvant, we sensitized mice twice with OVA and IMDQ+SDI or Alum and then switch the adjuvant and administered the mice two more sensitization doses. This was performed both in C57BL/6 mice, which typically present a Type 1-biased response, and BALB/c, that normally exhibit a Type 2-biased response, although this broad classification is an oversimplification, especially in the allergy context^55^. C57BL/6 showed high flexibility in their antibody subclass production and resulted a balanced Type 1/Type 2 response regardless of the adjuvant used for priming and the one use for boosting, as long as IMDQ+SDI was administered. While C57BL/6 primed with Alum produced OVA-binding IgG2c as soon as one week after they were boosted with IMDQ+SDI, BALB/c showed reduced ability to reshape their antibody response. In BALB/c, boosting with IMDQ+SDI did not lead to IgG2a production prior to challenge, however, after mice were challenge with OVA, they started producing detectable levels of IgG2a, which never reach the levels of animals primed with Alum. Due to the study design, we cannot conclude that the OVA challenge was the primary driver of the antibody class switching to IgG2a, as antibody production in BALB/c proved to be slower than in C57BL/6 and it is well within a reasonable timeline that those antibodies started to be produced before the challenge. While Type 2 to Type 1 flexibility in BALB/c was very limited, Type 1 to Type 2 flexibility was high and BALB/c mice primed with IMDQ+SDI caught up to the levels OVA-binding IgG1 in the Alum group, as soon as one week after the second Alum booster.

Antibody subclasses are a good proxy for immune response polarization; however, they might not be the main immune players involved in the prevention of allergy sensitization with IMDQ+SDI. To test this hypothesis, we performed IgG2b and IgG2c adoptive transfer purified from IMDQ+SDI sensitized mice, into C57BL/6 sensitized with OVA-Alum. After challenge we could observe that there were no differences between groups receiving the adoptive transfer and the group that was administered PBS. If anything, a non-significant increase in morbidity after each OVA challenge could be observed in the groups that received the adoptive transfer. This increased morbidity might be a sign of anaphylaxis after challenge, which is driven by IgG subclasses in mice. This phenomenon has been previously observed in mice sensitized with IgG1, IgG2a, or IgG2b anti-trinitrophenyl mAbs and challenge with trinitrophenyl-BSA intravenously^33^.

One of the hallmarks of the OVA-induced asthmatic allergy model, is the presence of elevated IgE after OVA challenge^19^, although levels vary depending on mice genetic background^56^. In naïve C57BL/6, we showed that sensitization with IMDQ or IMDQ+SDI reduced the levels of total IgE when compared to the Alum groups. Still, even in Alum groups, levels of IgE were limited. Elevating the number of sensitizations and challenges led to increased IgE levels in animals sensitized with Alum in the second experiment. Those levels were significantly reduced by either priming or boosting with IMDQ+SDI. Thus, we showed that even Alum-sensitized C57BL/6 could revert their allergic antibody response when boosted with IMDQ+SDI. Nevertheless, BALB/c mice primed or boosted with IMDQ+SDI, although they presented a balanced Type 1/Type 2 antibody response after challenge, showed no significant differences in IgE levels at an early timepoint after challenge (7 DPC). Still, IMDQ+SDI had a relevant effect on the IgE kinetics that wane down significantly faster than in the Alum group. IgE has been shown to steadily increase after OVA-challenge in OVA-sensitized mice until 28DPC and it is maintained for several more weeks^57,58^, suggesting a long-term effect of IMDQ+SDI sensitization in BALB/c.

Mucosal barriers are the surfaces where the first encounter with the allergen takes place. Generally, intramuscular vaccination induces poor mucosal immunity^59^. Still, we tested the production of OVA-binding IgG subtypes and IgA in the mucosal surfaces. In naïve C57BL/6 mice, we observed that only mice receiving IMDQ or IMDQ+SDI produced OVA-binding IgA. While the role of IgA remains unclear in allergic responses, clinical observations correlate low levels of secretory-IgA with elevated asthma, allergic symptoms and reduced lung function^60^. In mice, secretory-IgA has been linked to the downregulation of proinflammatory responses associated with the uptake of potentially allergenic antigens^61^. In previously Alum-sensitized C57BL/6 mice, boosting with IMDQ+SDI led to OVA-binding IgA production in BALF, that was absent in Alum sensitized mice. Interestingly, BALB/c mice sensitized only with Alum presented OVA-binding IgA in BALF, that was similar to Alum➔IMDQ+SDI animals but reduced compared to the levels in IMDQ+SDI➔Alum mice. BALB/c, but not C57BL/6, are known to have polyreactive IgAs^62^, which may impact the mucosal IgA response to the allergen. Levels of OVA-binding IgG subtypes in BALF, reflected the results observed in serum for both C57BL/6 and BALB/c.

Focusing on cytokine profiling in BALF and lung tissue, results also supported the Type 1/Type 2-biased immune response shaped by the different adjuvants. Naïve C57BL/6 sensitized mice presented elevated levels of IL-4 and IL-5 in BALF as soon as 1 DPC, as it is expected in this OVA allergic model^63^. At later timepoints (11DPC), other cytokines such as IL-9, IL-17A, IL-22 and IL-23 presented elevated levels, all of which have been previously linked to allergic airway inflammation in mice^64–67^. Elevating the number of Alum sensitizations and OVA challenges, resulted in increased levels of other cytokines in lung homogenates, including IL-13, IL-6, GM-CSF, IL-10 or eotaxin. IL-13 is expected to be induced in the Type 2 context, although its role in asthmatic allergy is still debated, as efforts in blocking IL-13 signaling has yielded no positive results in human trials and only simultaneous IL-4/IL-13 blockage has shown effective^68^. Eotaxin is a well-known eosinophil recruiting chemokine, normally elevated in models of allergic asthma and correlated with the increased eosinophil recruitment observed in Alum groups. However, its exogenous administration has shown no effect on airway hyperresponsiveness^69^. IL-6, IL-10 and GM-CSF are also known to regulate allergic airway hyperresponsiveness in mice^70–72^. Naïve C57BL/6 mice sensitized with IMDQ or IMDQ+SDI presented elevated levels in BALF of many classic Type 1 cytokines such as TNF-α, IL-1β, IFN-γ or IL-2, clearly defining a Type 1-polarizing cytokine/chemokine signaling in these animals.

In the Alum➔IMDQ+SDI and the IMDQ+SDI➔Alum groups, allergic phenotype was corrected or prevented, respectively. In both cases, IMDQ+SDI sensitization reduced hallmark Type 2 cytokines, such as IL-4, IL-5, IL-13. However, the reduction of these cytokines only reflected a decline in lung eosinophils in C57BL/6, highlighting the strain-specificity in eosinophil recruitment dynamics, which is true not only in allergy but also after infection^63,73^. For other Type 2 cytokines, such as IL-6 or IL-33, reduction in IMDQ+SDI sensitized groups was more relevant in C57BL/6 than BALB/c. IL-33 has been associated in C57BL/6 with a self-perpetuating amplification loop that maintains chronic airway inflammation^74^ while IL-6 has been linked to pulmonary eosinophilia in this model^75^. Additionally, IMDQ+SDI administration increased Th1-associated markers, particularly IL-1β, CXCL10, TNF-α, and IL-27. Once again increase in those cytokines was more obvious in C57BL/6. While lung IFN-γ levels increased modestly, the increase is further supported in the periphery by the ELISpot assays showing significantly elevated IFN-γ–producing splenocytes in both IMDQ+SDI C57BL/6 groups, but only in IMDQ+SDI➔Alum BALB/c. Chemokine CCL5 (also known as RANTES), also increased with IMDQ+SDI sensitization, along with CXCL2 and CCL4, but only in C57BL/6 and more dramatically in IMDQ+SDI➔Alum. Interestingly CCL5 treatment, has been shown to prevent allergic airway hyperresponsiveness, but only in the context of repeated allergen challenge, similar to our model^76^. All these changes suggest a shift toward a restrained Type 1-skewed lung environment, especially in C57BL/6 mice, without a concurrent rise in Th17-related markers, such as IL-17A or IL-23^77^, thus suggesting a protective role of the adjuvant, rather than am inflammatory Th17/neutrophilic phenotype.

MCIA analysis comparing Alum and Alum➔IMDQ+SDI revealed a pivotal role of IL-1β in the reprogramming allergic inflammation toward a protective phenotype, not only in the lung but also in the periphery (spleen). IL-1β has been classically associated with an allergy-promoting role^78^. However other models, such as early-life rhinovirus infection in mice which promotes asthma development later in life, suggest that IL-1β has a protective role by limiting Type 2 inflammation and mucous metaplasia, which could prevent the onset of asthma^79^. Interestingly, the increase in IL-1β was independent of IL-12p70. The increased levels of IL-1β could also be caused by a synergistic effect of both adjuvants, as Alum is known to activate the NLRP3 inflammasome promoting caspase-1-dependent IL-1β and IL-18 release^80^.

During the 2009 influenza pandemic, asthma was found to be the most common comorbidity among patients hospitalized with influenza^81^. However, the role of asthmatic allergy in the outcome of influenza infections is still poorly understood^23^ and studies have presented conflicting evidence. Various reports suggest that allergic asthma contributes to accelerated clearance of influenza and influenza resistance due to DCs and NKs priming by the allergen, as well as the prevention of immunopathology^25,82,83^. Other authors suggest an increased susceptibility to heterologous influenza secondary infection in allergy sensitized mice^84^. We have shown before that IMDQ+SDI enhances Type 1-biased influenza vaccine responses^14^, and we hypothesized that priming animals with IMDQ+SDI, even in combination with an unrelated antigen, could improve immune responses against a subsequent influenza infection. Surprisingly, although results after OVA-challenge were strikingly different in BALB/c and C57BL/6, sensitization with IMDQ+SDI (regardless of its use for prime or boost) led significantly reduced morbidity and lung viral titers of sublethal influenza infection compared to the Alum-sensitized group and a similar morbidity to the PBS group in both mouse strains. This suggest that Alum sensitization increase influenza infection morbidity and viral titers rather than IMDQ+SDI improving the outcome of the infection compared to the PBS group. Alum-sensitized mice, presented increased percentages of T_regs_ in the lung at 5DPI. It has been suggested that distinct waves of T_regs_ are recruited to the lung after influenza infection, with a first wave expressing a strong interferon-stimulated gene signature that appeared at 3DPI, peaking at 7DPI, and waning down by 21DPI. A second diverse wave of tissue-repair-like T_regs_ starts at 10DPI and is then maintained through 21DPI^85^. It is possible, that in a lung environment of the Alum-sensitized mice were cytokines like IL-4,5,10 and 13 were elevated, early recruitment of tissue-repair-like T_regs_ was favored, which would not promote viral clearance. Additionally, Alum groups presented an enhanced depletion of alveolar macrophages that were replaced by both iNOS^+^ and ArgI^+^ monocyte-derived macrophages. Alum-sensitized groups also presented increased activation of both neutrophils and eosinophils (measured via SiglecF expression). SiglecF^hi^ neutrophils are known to be generated in the context of lung damage caused by air-pollutant and lead to the exacerbation of airway inflammation^86^. SiglecF^hi^ eosinophils have been linked to protection in breakthrough influenza infections^27^, however in the absence of a Type 2 cytokine profile, which it is present in the Alum-sensitized groups.

Finally, we evaluated the role of T-cells in the Type 1-biased OVA response in IMDQ+SDI sensitized mice. We could observe that loss of CD4^+^ T-cells abrogated the reduction of lung eosinophil recruitment, suggesting that a subpopulation of CD4^+^ T-cells is restraining eosinophil recruitment in the OVA-IMDQ+SDI mice. This result conflicts with the notion that CD4^+^, but not CD8^+^ T-cells, mediate antigen-induced eosinophil recruitment in the airways in OVA-sensitized models^87^. The modified lung environment by IMDQ+SDI might play a role on this different CD4^+^ function, as IL-5 mediates the eosinophil recruitment, and it is reduced in our model. On the other hand, the CD4^+^CD25^+^ T-cell subset has been identified as a key modulator of Th2-mediated pulmonary inflammation via the suppression of the development of a Th2-phenotype, highly effective at promoting airway eosinophilia^88^. CD8^+^ T-cells seem to govern the eosinophil subtype/activation status, as SiglecF^hi^ eosinophils dramatically increase in CD8^+^-depleted mice. Strikingly, simultaneous CD4^+^/CD8^+^-depletion does not have a similar effect on this eosinophil subpopulation, suggesting that CD4^+^ T-cells are needed to recruit sufficient eosinophil numbers for CD8^+^ T-cells to shift the lung microenvironment and promote eosinophil activation. In a similar fashion CD8^+^ T-cell depletion affected neutrophil maturity, but the effect was not observed when the two T-cell subsets were depleted.

Conclusively, we demonstrated in this manuscript the OVA-immunization with IMDQ+SDI prevents allergic sensitization via the induction of a balance Type 1/Type 2 response. The adjuvant achieved the reversion of the allergic phenotype even in mice previously sensitized with OVA-Alum. However, the prevention of the allergic phenotype was dependent on genetic background. Only partial re-shaping of the immune response was achieved in BALB/c mice, and became more apparent at later timepoints, while early allergic responses were not strongly modified. Still, both mouse strain prime or boosted with IMDQ+SDI presented a reduced morbidity of a secondary influenza challenge when compared to the Alum group. Finally, we demonstrated that IgG2c, by itself, cannot protect from allergy sensitization and that both CD4^+^ and CD8^+^ for IMDQ+SDI to prevent eosinophil recruitment and activation upon OVA-challenge.

## METHODS

### Ethics statement

All experiments were approved and carried out in compliance with the Institutional Biosafety Committee (IBC) and Institutional Animal Care and Use Committee (IACUC) regulations of Icahn School of Medicine at Mt. Sinai (New York, USA). Protocol numbers: 202100007 and 202400000024.

### Cells

Madin-Darby canine kidney (MDCK) cell line was maintained in Dulbecco’s Modified Eagle Medium (DMEM, Corning) supplemented with 10% Fetal bovine serum (FBS, HyClone) and 1X penicillin/ streptomycin (Pen-Strep, Corning).

### Virus

H1N1 A/New Caledonia/20/1999 (NC99) was grown in 8-days old embryonated chicken eggs and was titrated by plaque assay on pre-seeded MDCK cells. The half-lethal dose of the virus (LD_50_) was calculated based on intranasal infections in 6-8 weeks old naïve female C5BL/6 and BALB/c (The Jackson Laboratory) mice using 10-fold serial dilutions of the virus. LD_50_ in BALB/c was established at 3064 PFU (PFU, plaque forming units) and 1532 PFU in C57BL/6.

### OVA

Imject™ ovalbumin (OVA, Thermo Scientific) was diluted to a stock concentration of 10 mg/mL in PBS, aliquoted and stored at -20°C until use. Sensitizations and challenges were performed with 20 µg of OVA per mice.

### Adjuvants

Alhydrogel adjuvant 2% (Alum, InvivoGen) was used for OVA sensitizations either intraperitoneally or intramuscularly. Right before administration, Alum was vigorously mixed.

The SDI RNA was transcribed in vitro using AmpliScribe T7 high yield transcription kit from Lucigen according to manufacturer’s instructions. Quality controls were performed as described before^14^. 1 μg per mouse was used for each sensitization.

IMDQ-PEG-CHOL (IMDQ, for short) was synthesized as previously described^17^. IMDQ adjuvant was mixed with OVA at an equivalent of 10 μg core IMDQ (100 μg of IMDQ-PEG-CHOL) per mouse. All intraperitoneal injections were performed in a final volume of 100 μL while intramuscular administrations were performed in 50 μL. PBS was used to dilute OVA-adjuvant mixes to the final injection volumes and they were mixed by vortexing during 30sec right before injection.

### Study design

In a first experiment, forty-six 6-8 week-old naïve female C5BL/6 mice were sensitized twice, one week apart, with either: (1) 50 µL of PBS intramuscularly; (2) 20 µg of OVA adsorbed to 2% Alum intraperitoneally (OVA-Alum IP); (3) 20 µg of OVA adsorbed to 2% Alum intramuscularly (OVA-Alum IM); (4) 20 µg of OVA combined with 10 μg of IMDQ or (5) 20 µg of OVA combined with 10 μg of IMDQ and 1 µg of SDI (OVA-IMDQ+SDI). One week after each sensitization, blood was drawn from each mouse for serum collection. One week after the second sensitization, mice were challenged intranasally under mild ketamine/xylazine anesthesia with 20 µg of OVA diluted in PBS into a final volume of 50 µL. After the challenge, mice body weights were monitored daily until 7 DPC (DPC, days post-challenge). One day after the OVA challenge (1 DPC), 4-5 mice per group were euthanized, mice were bled and bronchoalveolar lavage fluid (BALF) was collected. The remaining mice were euthanized at 11 DPC and the same tissues described for 1 DPC were collected (Fig. 1A).

In a second experiment, forty-eight 6-8 week-old naïve female C5BL/6 mice and forty-eight 6-8 week-old naïve female BALB/c mice were divided into 4 different groups (each mouse strain). All groups were sensitized intramuscularly four times over a period of four weeks with different adjuvants: (1) PBS four times, (2) OVA-Alum four times, (3) OVA-Alum the first two sensitizations and then OVA-IMDQ+SDI the last two, (4) OVA-IMDQ+SDI the first two sensitizations, then OVA-Alum the last two. Mice were bled weekly for serum collection during the sensitization period. One week after the last sensitization mice were challenged intranasally three times over a period of five days with 20 µg of OVA each challenge, diluted to a final volume of 30 µL in PBS (challenged on 0, 2 and 4 DPC; rested on 1 and 3 DPC). Mice body weights were monitored daily until 7 DPC. One day after the last challenge (5 DPC) 4 mice per group were euthanized and lungs, spleens and blood were collected. 7 DPC, 4 additional mice per group were euthanized and BALF, spleens and blood were collected. The remaining mice were bled weekly for serum collection until 30 DPC. At this time, all remaining mice were infected intranasally with a sublethal dose (0.2 LD_50_) of NC99 diluted in PBS into a final volume of 50 µL. Mice body weights were monitored daily until 5 days post-infection (5 DPI). On 5 DPI, mice were euthanized, and lungs and blood were collected (Fig 2A).

In a third experiment we attempted to characterize the role of OVA-binding IgG2 antibodies in the immune response observed in the previous sections. With that aim, fifteen 6–8-week-old naïve female C5BL/6 mice were sensitized twice with OVA-Alum following the schedule described for the first experiment. Prior to challenge, mice were divided in 3 groups (n=5): (1) Control group injected with PBS intraperitoneally 3hr before OVA-challenge (OVA-Alum), (2) Group that received an adoptive transfer of IgG2c intraperitoneally, purified from pooled serum of mice sensitized with OVA-IMDQ+SDI in previous experiments, 3hr before OVA-challenge (OVA-Alum + IgG2c) and (3) Adoptive transfer of IgG2b intraperitoneally, purified from pooled serum of mice sensitized with OVA-IMDQ+SDI in previous experiments, 3hr before OVA-challenge (OVA-Alum + IgG2b). All mice were challenged intranasally three times over a period of five days with 20 µg of OVA each challenge, diluted to a final volume of 30 µL in PBS. Mice body weights were monitored daily until 5 DPC. One day after the last challenge (5 DPC) mice were euthanized and lungs and blood were collected (Fig. S11A).

In a final experiment, we set out to characterize the role of CD4^+^ and CD8^+^ T-cells in the Th1-polarization of the immune response to OVA caused by IMDQ+SDI. For that, twenty 6– 8-week-old naïve female C5BL/6 mice were sensitized with OVA-IMDQ+SDI twice following the experimental layout described for the first experiment. Prior to challenge, mice were then divided into four groups: (1) Control (OVA-IMDQ-SDI); (2) CD8^+^ T-cells were depleted three and one day prior to challenge (OVA-IMDQ-SDI CD8^+^ Depletion), (3) CD4^+^ T-cells were depleted three and one day prior to challenge (OVA-IMDQ-SDI CD4^+^ Depletion) and (4) Both CD8^+^ and CD4^+^ T-cells were depleted three and one days prior to challenge (OVA-IMDQ-SDI CD4/8 Depletion). All mice were challenged intranasally three times over a period of five days with 20 µg of OVA each challenge. Body weights were monitored daily until 5 DPC when mice were euthanized, and lungs and blood were collected (Fig. 7A).

### Serum collection

For non-terminal sample collections, blood was obtained via the submandibular route^89^, while in terminal procedures cardiac bleeding was performed. In all cases, blood was allowed to coagulate at 4°C overnight. Coagulated blood was then centrifuged at 400 x g for 5 minutes at 4°C and serum was collected and stored at -20°C until further analyses.

### Bronchoalveolar lavage fluid collection

Bronchoalveolar lavage fluid (BALF) was collected with a 18G catheter into 1.2 mL PBS with 2 mM EDTA (PBS-EDTA). Fluid was collected by introducing 0.6 mL of PBS-EDTA into the lungs through the trachea, gently massaging the thoracic cavity and retrieving the PBS-EDTA with the syringe. This process was repeated twice to a final volume of 1.2 mL.

### Enzyme-linked immunosorbent assay (ELISA)

To measure OVA-specific total IgG, IgG1, IgG2a, IgG2b or IgG2c Nunc MaxiSorp 96-well plates (ThermoFisher) were coated with 50 µL/well OVA to a final concentration of 500 ng/well in carbonate-bicarbonate buffer and incubated overnight at 4°C. Plates were washed 3 times with PBS-T then blocked with 100 µL/well of blocking buffer (5% milk in PBS-T) and incubated for 1 h at room temperature (RT). During the blocking step, serum samples were serially diluted 3-fold a total of 7 times, with a starting dilution of 1:50 1:100 or 1:150, depending on the experiment. After blocking, 50 µL/well of the serial dilutions was added to plates and incubated for 1.5 hr at RT. Plates were washed 3 times with PBS-T then 100 µL/well of goat anti-mouse IgG-HRP (ab6823, Abcam) diluted 1:5000 in blocking buffer or goat anti-mouse IgG1-HRP (1071-05, SouthernBiotech), goat anti-mouse IgG2a-HRP (1081-05, SouthernBiotech), goat anti-mouse IgG2b-HRP (1091-05, SouthernBiotech), goat anti-mouse IgG2c-HRP (1078-05, SoutherBiotech) diluted 1:4000 in blocking buffer was added to each well and incubated for 1 hr at RT, depending on the specific ELISA. After washing plates 3 times with PBS-T, 50 µL/well of 1-Step Turbo TMB Substrate (ThermoFisher) was added and then incubated for 20 minutes at RT. The reaction was stopped by adding 50 µL/well of ELISA Stop Solution (Invitrogen).

To measure total IgE, plates were coated with 100 µg/well of anti-mouse IgE antibody (Clone R35-72, BD). The protocol was analogous to the ones described before. As a secondary antibody, 100 µL/well of goat anti-mouse IgE-HRP (1110-05, SouthernBiotech) were used.

To evaluate mucosal antibody response, OVA-specific total IgG, IgG1, IgG2a, and IgG2c were measured as described above, with a 1:10 starting dilution of BALF. OVA-specific IgA was measured in analogously with goat anti-mouse IgA-HRP (1040-05, SouthernBiotech) diluted 1:4000 in blocking buffer To determine specific antibody responses against NC99, an analogous protocol was followed, with plates coated with 50 µL/well trivalent inactivated influenza virus vaccine (TIV; Fluzone® Influenza Virus Vaccine containing the matched influenza A H1N1 component (A/New Caledonia/20/1999/IVR-116) diluted 1:200 in carbonate-bicarbonate buffer.

In all cases, plates were read at 450 nm and 650 nm. Background subtracted optical density values (OD450-650nm) were used for downstream analyses. The average value of the negative controls plus three times the standard deviation was established as a cut-off for positive results. Area under the curve or endpoint-titer calculations were performed for representation with GraphPad Prism.

### Flow cytometry

Either BALF or lung tissues were used for the profiling of the cell populations in the alveolar space and lung tissue via flow cytometry. When working with lung tissue, left lung lobes were collected and single cell suspensions were generated using the Mouse Lung Dissociation kit from Miltenyi Biotech following manufacturer’s instructions. In brief, 2.4 mL of 1× Buffer S were added to a gentleMACS C Tube and kept on ice until lung lobes were harvested. Then 100 µL of Enzyme D and 15 µL of Enzyme A added. C Tubes were tightly closed and attached upside down onto the sleeve of the gentleMACS Octo Dissociator and program 37C_m_LDK_1 was performed.

After generating a single cell suspension either from lung tissue or BALF, cells were centrifuged for 5 mins at 500 x g and the supernatant was decanted. Then, cells were resuspended into 2 mL of RBC lysis solution and incubated at RT for 10 min. Then 10 mL of PBS was added to stop lysis and cells were centrifuged for 5 mins at 450 x g and the supernatant was decanted. Then cells were resuspended into 50 μL of Fc Block and incubated for 5 mins at RT. Meanwhile the different staining cocktails were prepared. Then several flow cytometry panels were used in the different experiments.

One panel was used for the broad characterization of myeloid lung cells performed in the first and second experiments. With that purpose, the following antibodies volumes were diluted into a final 50 μL antibody cocktail in FACS buffer for surface staining: 1 μL of BD Pharmingen™ FITC Hamster Anti Mouse CD3e (Clone 145-2C11), BD Pharmingen™ PE Rat Anti-CD11b (Clone M1/70), BD Horizon™ PE-CF594 Rat Anti-Mouse Siglec-F (Clone E50-2440), BD Pharmingen™ PE-Cy7 Hamster Anti-Mouse CD11c (Clone HL3), Ly-6C Monoclonal Antibody, APC, eBioscience™, Invitrogen™ (Clone HK1.4), Alexa Fluor® 700 anti-mouse Ly-6G Antibody (Clone 1A8), 0.5 μL of Fixable Viability-e780 and MHC Class II (I-A/I-E) Monoclonal Antibody, eFluor™ 450, eBioscience™, (Clone M5/114.15.2). The staining cocktail was added on top of the samples with the Fc Block, yielding a final volume of 100 μL, and they were incubated for 20 mins at RT protected from the light. After the incubation period, cells were centrifuged for 5 mins at 500 x g and washed twice with 200 μL of FACS buffer, and passed through a cell strainer snap cap tube (Falcon) right before acquiring the samples in the cytometer. Samples were acquired using either a Beckman Coulter Gallios flow cytometer (first experiment) or an Attune NxT flow cytometer (second experiment).

After the NC99 challenge, further assessment of monocyte-macrophage subpopulations and T-cell subpopulations in the lung was performed. For that, lung tissue was processed as described before and two panels were used. For the T-cell subpopulations, the following surface staining was performed: 2 μL of BV480 CD3 anti-mouse monoclonal antibody (Clone 17A2, eBioscience); 1 μL of eFluor 450 CD25 rat anti-mouse monoclonal antibody (Clone PC61.5, eBioscience) and PE/Dazzle™ 594 anti-mouse CD192 (CCR2) antibody (Clone SA203G11, BioLegend); 0.63 μL of AF532 anti-mouse CD45.2 Monoclonal Antibody (Clone 104, eBioscience™, Invitrogen™); 0.5 μL of Brilliant Violet 605™ anti-mouse CD4 Antibody (Clone RM4-5, BioLegend) and BD Horizon™ Fixable Viability Stain AF520 (BD Bioscience); PerCP anti-mouse CD8a Antibody (Clone 53-6.7, BioLegend).

For the monocyte-macrophage characterization the following antibodies were used: surface staining: 1 μL of eFluor™ 450 Anti-mouse CD11c Monoclonal Antibody (Clone N418, eBioscience™, Invitrogen™), PE/Dazzle™ 594 anti-mouse CD192 (CCR2) Antibody (Clone SA203G11, BioLegend) and PerCP-eFluor™ 710 anti-mouse MHC Class II (I-A/I-E) Monoclonal Antibody (Clone M5/114.15.2, eBioscience™, Invitrogen™); 0.63 μL of AF532 anti-mouse CD45.2 Monoclonal Antibody (Clone 104, eBioscience™, Invitrogen™), BV510 anti-mouse Ly-6C Antibody (Clone HK1.4, BioLegend), BV650 anti-mouse F4/80 Monoclonal Antibody (Clone BM8, eBioscience™); 0.5 μL of BD OptiBuild™ BV786 Rat Anti-Mouse Siglec-F (Clone E50-2440, BD Biosciences), BV480 anti-mouse Ly-6G Monoclonal Antibody (Clone 1A8-Ly6g, eBioscience™); 0.25 μL of BV57 anti-mouse CD11b Antibody (Clone M1/70, BioLegend) and BD Horizon™ Fixable Viability Stain 575V (BD Bioscience). After the surface staining, samples were washed twice with FACS buffer and permeabilized for 30min at 4°C. Then, samples were washed twice and incubated for 15 min in 50 μL Fc block at RT. Then 50 μL of intracellular staining antibody cocktail was added on top and incubated for 20 min at RT, containing: PE-Cyanine7 iNOS anti-mouse Monoclonal Antibody (Clone CXNFT, eBioscience), PE Arginase I anti-mouse monoclonal antibody (Clone A1exF5, eBioscience). After the intracellular staining, samples were washed twice with FACS buffer and resuspended to a final volume of 150 μL. Samples stained with either of the two panels were kept overnight at 4°C and analyzed the following day in a Cytek® Northern Lights™ full spectrum flow cytometer.

Finally, in the IgG2 adoptive transfer and T-cell depletion experiments we focused again on myeloid lung cell populations, with a focus on eosinophil and neutrophil maturation and activation. For that, lung processing was performed as described above. The staining panel was: 2.5 μL of PerCP-Cy5.5 anti-mouse Ly6G (Clone 1A8, BD Bioscience); 1.25 μL of PerCP anti-mouse TER-119 (Clone TER-119, BioLegend); 0.625 μL of BV421 anti-mouse Siglec-F (Clone E50-2440, BD Bioscience), BV510 anti-mouse MHCII (Clone M5/114.15.2, BioLegend), FITC anti-mouse CD11c (Clone HL3, BD Bioscience), PE anti-mouse CD101 (Clone Moushi101, eBioscience), RB780 anti-mouse CD125 (Clone T21, BD Bioscience); 0.3125 μL BV650 anti-mouse CD11b (Clone M1/70, BioLegend); and 0.0625 μL of eFluor 450 Viability Dye (eBioscience). After staining for 45 min at 4°C, samples were washed and fixed with 1% PFA for 15 min at 4°C. After washing, samples were run the next day in a Cytek® Northern Lights™ full spectrum flow cytometer.

Data analysis was performed using FlowJo (Treestar) and compensated using the built-in AutoSpill algorithm. Data were visualized using Graphpad Prism 10.0.

### Measurement of cytokines and chemokines

Right middle and caudal lung lobes were collected and homogenized. The Cytokine & Chemokine 26-Plex Mouse ProcartaPlex™ Panel 1 (ThermoFisher) to measure cytokine and chemokine concentrations in the lungs, sometimes combined with Simplex IL-33. The assay was performed according to the manufacturer’s instructions and the plate was placed on an orbital shaker set to 300 rpm for all incubation steps. Briefly, 50 µL of lung homogenates were combined with beads in an optical bottom black 96-well plate and incubated for 30 minutes at RT protected from light. After 30 minutes, the plate was moved to 4°C for overnight incubation. The next day, the plate was equilibrated to RT for 30 minutes, then washed 3 times with 150 µL/well of 1X Wash Buffer diluted according to manufacturer’s instructions. Then, 25 µL/well of 1X Detection Antibody mixture was added and incubated at RT for 30 minutes. The plate was again washed and then 50 µL/well of 1X Streptavidin-PE solution was added and incubated for 30 minutes at RT. After once again washing, 120 µL/well of Reading Buffer was added and the plate was incubated for 5 minutes at RT. Subsequently, data were acquired on a Luminex 100/200 analyzer (Millipore) with xPONENT software (version 4.3). Data visualization and analysis were conducted using GraphPad Prism and R computing language in RStudio.

### IFN-**_γ_** and IL-4 ELISPOTs

Spleens were harvested from mice and collected in RPMI-1640 media supplemented with 10% FBS and 1x Penicillin/Streptomycin. Single cell splenocyte suspensions were prepared by forcing the spleens through a 70 μm cell strainer. Mouse IFN-γ^+^ (EL485, R&D Systems) and IL-4^+^ ELISPOTs (EL404, R&D Systems) assays were performed using 10^5^ cells/well following manufacturer’s instructions. In brief, all wells in the ELISPOT plate were incubated with 200 μL of sterile culture media for 20 min at RT. Then, media was aspirated and 10^5^ splenocytes were added per well in 100 μL of medium and splenocytes were stimulated overnight at 37°C and 5% CO_2_ with Ovalbumin or Hemagglutinin overlapping peptide pools (Miltenyi Biotec: PepTivator Ovalbumin, PepTivator Influenza A (H1N1) HA, respectively) or kept unstimulated. After incubation, plates were washed four times and then 100 μL of detection antibody mixture into each well, and incubated for 2 hours at RT on a rocking shaker. After four more washes, 100 μL of Streptavidin-AP was added into each well, and incubate for 2 hr at RT. After another four washes, 100 μL of BCIP/NBT substrate was added into each well, and incubated for 1 hr at RT protected from the light. Finally, the plates were thoroughly washed under tap water 5 times and allowed to air dry for 2 hr in the dark. Images of the plates were taken with a CTL Immunospot Analyzer and then images of each individual well were analyzed with Image J. The number of spots in each well were represented as the number of IFN-γ or IL-4 producing cells per million splenocytes.

### Determination of lung viral titers

Virus titration was performed by plaque assays from lung homogenates. In brief, the homogenates were 10-fold serially diluted in PBS and added on top of pre-seeded and PBS-washed MDCK monolayers on a 12-well plate. The cells were then incubated for 1 hr at RT, gently shaken every 10 min. The diluted samples were then removed, and the monolayers were washed with PBS. Lastly, 1 mL of the overlay mixture (2% oxoid agar and 2X minimal essential medium (MEM) supplemented with 1% diethyl-aminoethyl (DEAE)-dextran and 1 mg/ml tosylamide-2-phenylethyl chloromethyl ketone (TPCK)-treated trypsin) was added on top of the monolayers and incubated for 48 hr at 37°C, 5% CO_2_. After the incubation, plates were fixed in 4% formaldehyde and stored overnight at 4°C. The next day, the overlay was removed, and the plaques were immune-stained with IVR-180-post challenge polyclonal serum, diluted 1:1000 in blocking buffer. The plates were washed and incubated with goat anti-mouse IgG-HRP (ab6823, Abcam) diluted 1:5000 for 1h at RT with gentle shaking. After that, the plates were washed with water and the plaques were stained with True Blue substrate for 20 min. Finally the number of plaques were counted and represented as PFU/mL.

### IgG subtype purification

Purification of IgG1, IgG2b and IgG2c from C57BL/6 mice was performed as previously described^90^. Briefly, 2 mL of 0.1 M sodium phosphate (pH 8.0) were added to 4 mL of serum and then the mix was adjusted to pH 8.1 with 1M Tris-HCl (pH 9.0). In a ÄKTA start FPLC (Cytiva), we coupled a 1 mL Protein-A-Sepharose column (Cytiva) and pre-equilibrate both the FPLC pumps and the column with 0.1 M sodium phosphate (pH 8.0). Then we applied the sample and was the column with 25-30 mL of 0.1 M sodium phosphate (pH 8.0). Then, elution of IgG subtypes started with a 0.1M citrate buffer (pH=6) to obtain IgG1, followed by 0.1M citrate buffer (pH=4.5) to elute IgG2c and lastly 0.1M citrate buffer (pH=3.5) to obtain IgG2b. Each buffer was passed through the column in all elutions until absorbance readouts fell back to baseline. IgG2c and IgG2b fractions were passed one additional run through the column to ensure high purity levels. Right after elution neutralization of the elution buffer was performed with 1M Tris-HCl (pH 9.0) to pH 7.4. All buffers and samples were maintained at 4°C to minimize IgG functionality loss.

Purified fractions were passed through a 50kDa Amicon (Millipore) to exchange solution buffers for PBS. Right before injection in the mice, IgG2c and IgG2b solutions were forced through a 0.2 μm filter (Agilent) with a 5 mL syringe and 150 μL were used for intraperitoneal injection of the mice.

### T-cell depletion

For the depletion of the different T-cell subpopulations, mice were injected intraperitoneally with either CD4 depleting antibody BioXCell #BP0003-1 (clone YTS 177) with a dosing of 500 μg in 100 μL PBS, CD8 depleting antibody BioXCell #BE0223 (clone 53-5.8) with a dosing 250 μg in 100μL PBS or a combination of both, 72 and 24 hr prior to the first OVA challenge. Success of the depletion was evaluated by flow cytometry with blood samples the one day after the last OVA challenge (5 DPC).

### Multiple co-inertia (MCIA) analysis

To model how antibody, cytokine and cellular features combine to define differential immune skewing by adjuvant combinations within and across mouse strains, the analytical method of Multiple co-inertia analysis (MCIA) was deployed in R. This method uses machine learning algorithms to combine distinct Omics into a single multi-Omic dataframe that allows for more direct comparison of the relationship between features from different Omics with each other and with an outcome of interest (i.e., adjuvant treatment, mouse strain, etc). Synthetic variables (SynVars), similar to principle components in a traditional PCA, are generated that identify the largest drivers of variance within the multi-Omic dataset. These SynVars can then be used to explore differences between experimental conditions to identify groups of features which drive distinct immunological states. Further refinement of MCIA results for network modeling was performed by identifying SynVars that were significantly different comparing groups of interest and then identifying the individual features within those SynVars that were significantly associated to one or more outcome groups. Using CytoScape, network models were generated showing connections between key antibody, cytokine and cellular features with adjuvant combinations and mouse strains to highlight the key drivers of differences. This approach unifies the findings of the individual Omic analyses and highlights the key features that function together to define differential immune states induced by combinations of adjuavnts within and across mouse strains.

### Other statistical analysis

Other statistical analyses were performed with Graphpad. Unless otherwise specified, OVA-Alum group was established as control for multiple comparisons analysis that were performed either by Kruskal-Wallis one-way ANOVA with Dunn’s multiple comparisons test or two-way ANOVA with Tukey’s multiple comparisons test, the latter when several time points were analyzed. In all cases, significance is represented as: *p = 0.05 to 0.01, **p = 0.01 to 0.001, ***p = 0.001 to 0.0001, ***p < 0.0001.

## ACKNOWLEDGEMENTS

Research in the M.S. laboratory is funded by NIH/NIAID grants R21AI180874, R21AI176069, R01AI160706, R01DK130425, and partly funded by CRIPT (Center for Research on Influenza Pathogenesis and Transmission), a NIH NIAID-funded Center of Excellence for Influenza Research and Response (CEIRR, contract number 75N93021C00014).

