## Supplementary files for "A combination TLR7/8 and RIG-I agonist adjuvant reverts asthmatic allergic sensitization and prevents aggravated influenza infection in OVA-sensitized mice"

**TITLE:**

**SUPPLEMENTARY FIGURES**

**Supplementary Fig. 1.** BALF cytokine/chemokine profile in OVA sensitized C57BL/6 with different adjuvants one day after challenge.

**Supplementary Fig. 2.** BALF cytokine/chemokine profile in OVA sensitized C57BL/6 with different adjuvants eleven day after challenge.

**Supplementary Fig. 3.** BALF selected immune cell populations in OVA sensitized C57BL/6 with different adjuvants eleven day after challenge.

**Supplementary Fig. 4.** Lung cytokine/chemokine profile in C57BL/6 primed with Alum or IMDQ+SDI and boosted with the other adjuvant.

**Supplementary Fig. 5.** Lung cytokine/chemokine profile in BALB/c primed with Alum or IMDQ+SDI and boosted with the other adjuvant.

**Supplementary Fig. 6.** Spleen cytokine/chemokine profile in C57BL/6 and BALB/c primed with Alum or IMDQ+SDI and boosted with the other adjuvant.

**Supplementary Fig. 7.** OVA-specific and IgE antibody response after influenza challenge.

**Supplementary Fig 8.** Number of cells in the lung for different T-cell subsets at 5DPI after influenza challenge.

**Supplementary Figure 9.** Lung Cytokine/chemokine protein concentration at 5DPI after influenza challenge in C57BL/6

**Supplementary Figure 10.** Lung Cytokine/chemokine protein concentration at 5DPI after influenza challenge in BALB/c

**Supplementary Figure 11.** Multiple co-inertia analysis reveals similar effects on NC99 challenge outcome of OVA-sensitization in combination IMDQ+SDI in BALB/c and C57BL/6

**Supplementary Figure 12.** Purification of C57BL/6 IgG subclasses from serum using a Protein-A-sepharose column.

**Supplementary Figure 13.** Transfer of IgG2c or IgG2b has no protective effect on in Alum-sensitized mice

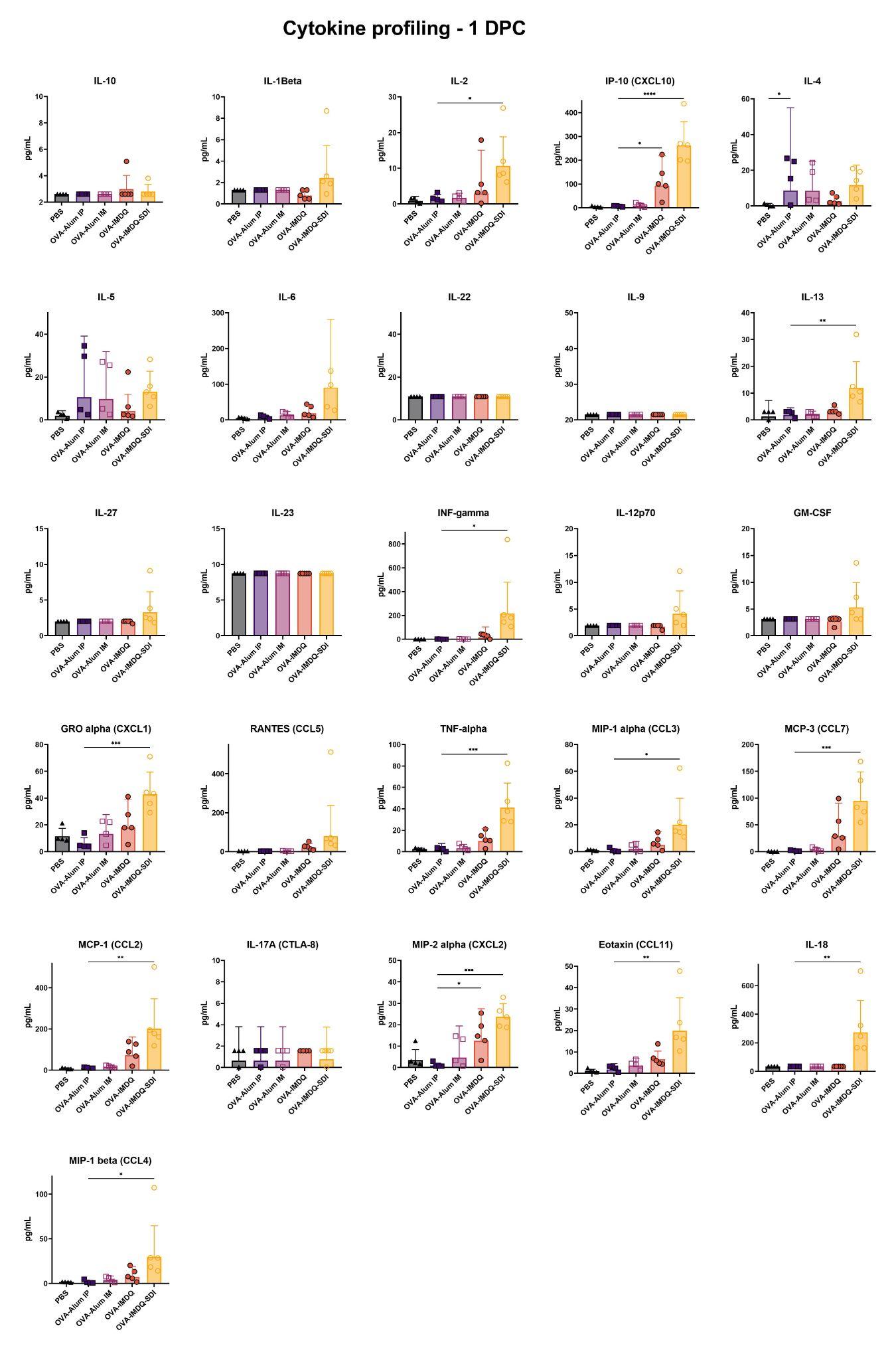

**Supplementary Fig. 1. BALF cytokine/chemokine profile in OVA sensitized C57BL/6 with different adjuvants one day after challenge.**

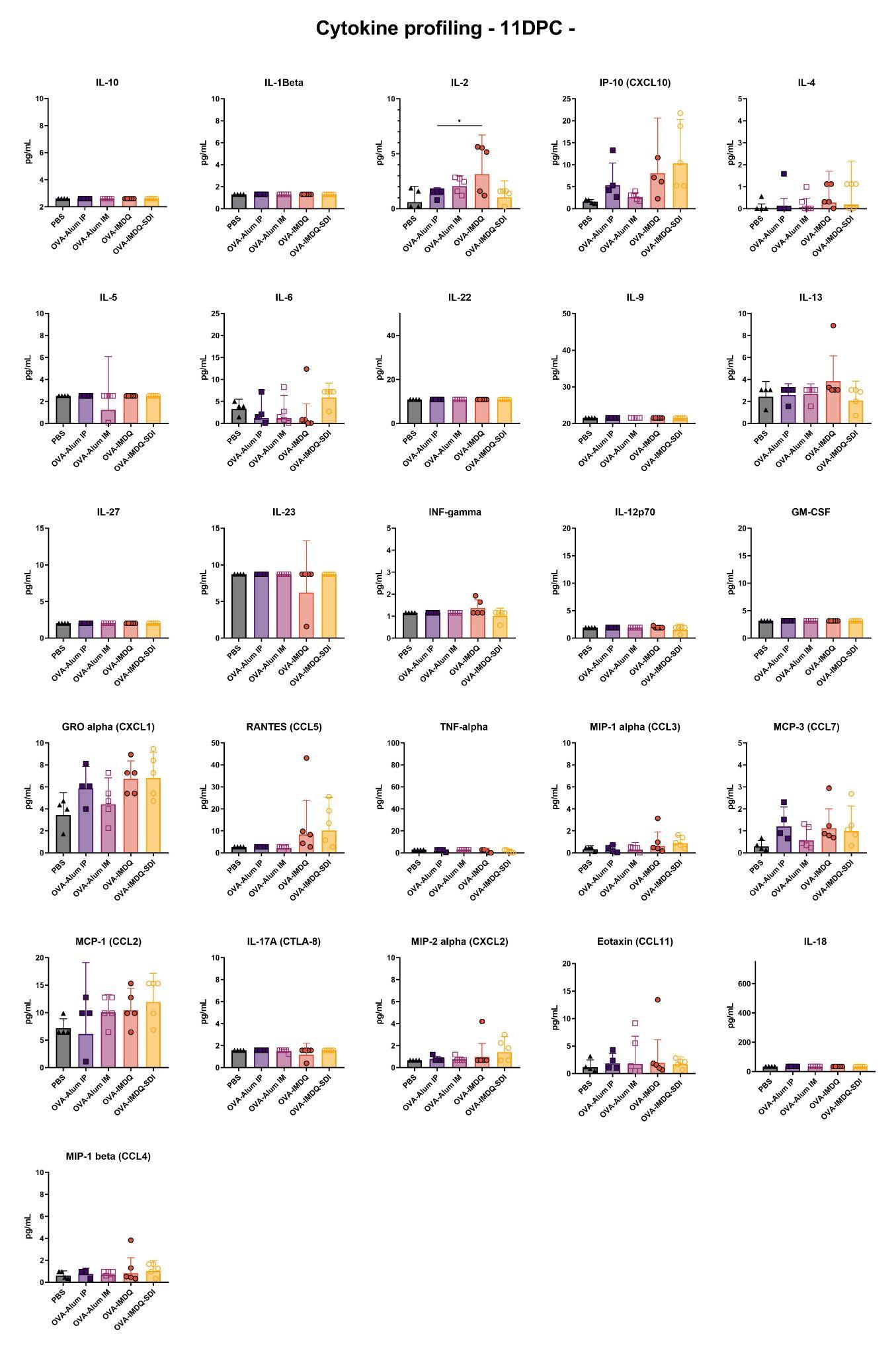

**Supplementary Fig. 2. BALF cytokine/chemokine profile in OVA sensitized C57BL/6 with different adjuvants eleven day after challenge.**

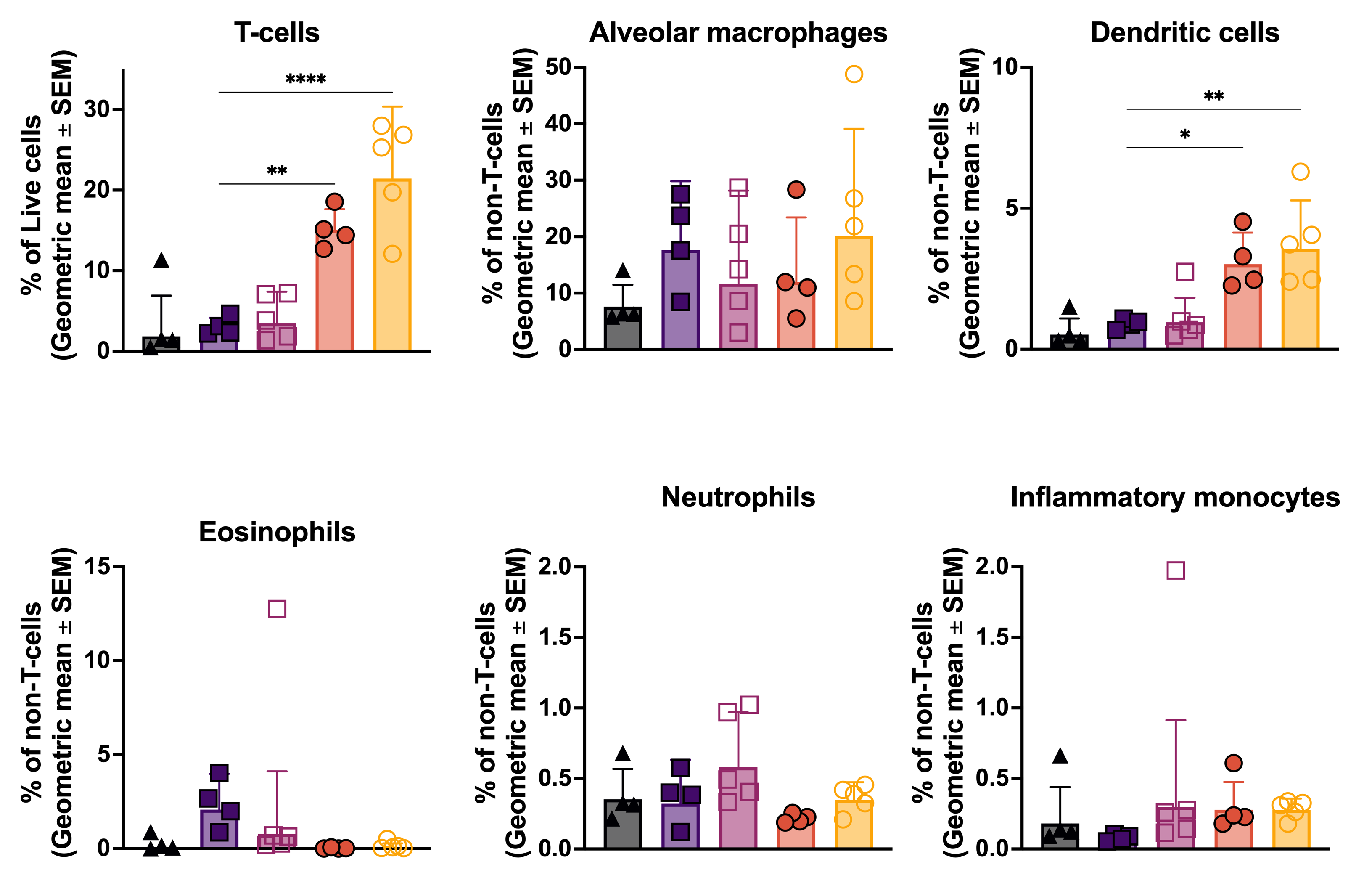

**Supplementary Fig. 3. BALF selected immune cell populations in OVA sensitized C57BL/6 with different adjuvants eleven day after challenge.** From left to right and top to bottom: T-cells (Gating: Live>CD3^+^), alveolar macrophages (Gating: Live > CD3^-^ > Ly6G^-^ > CD11c^+^ CD11b^-^ > SiglecF^+^), dendritic cells (Gating: Live > CD3^-^ > Ly6G^-^ > CD11c^+^>MHCII^+^ SiglecF^-^ ), eosinophils (Gating: Live > CD3^-^ > Ly6G^-^ > CD11c^-^ CD11b^+^ > SiglecF^+^), neutrophils (Gating: Live > CD3^-^ > Ly6G^+^ CD11b^+^ ), Inflammatory monocytes (Gating: Live>CD3^-^> Ly6G^-^ > CD11c^-^ CD11b^-^ > Ly6C^+^).

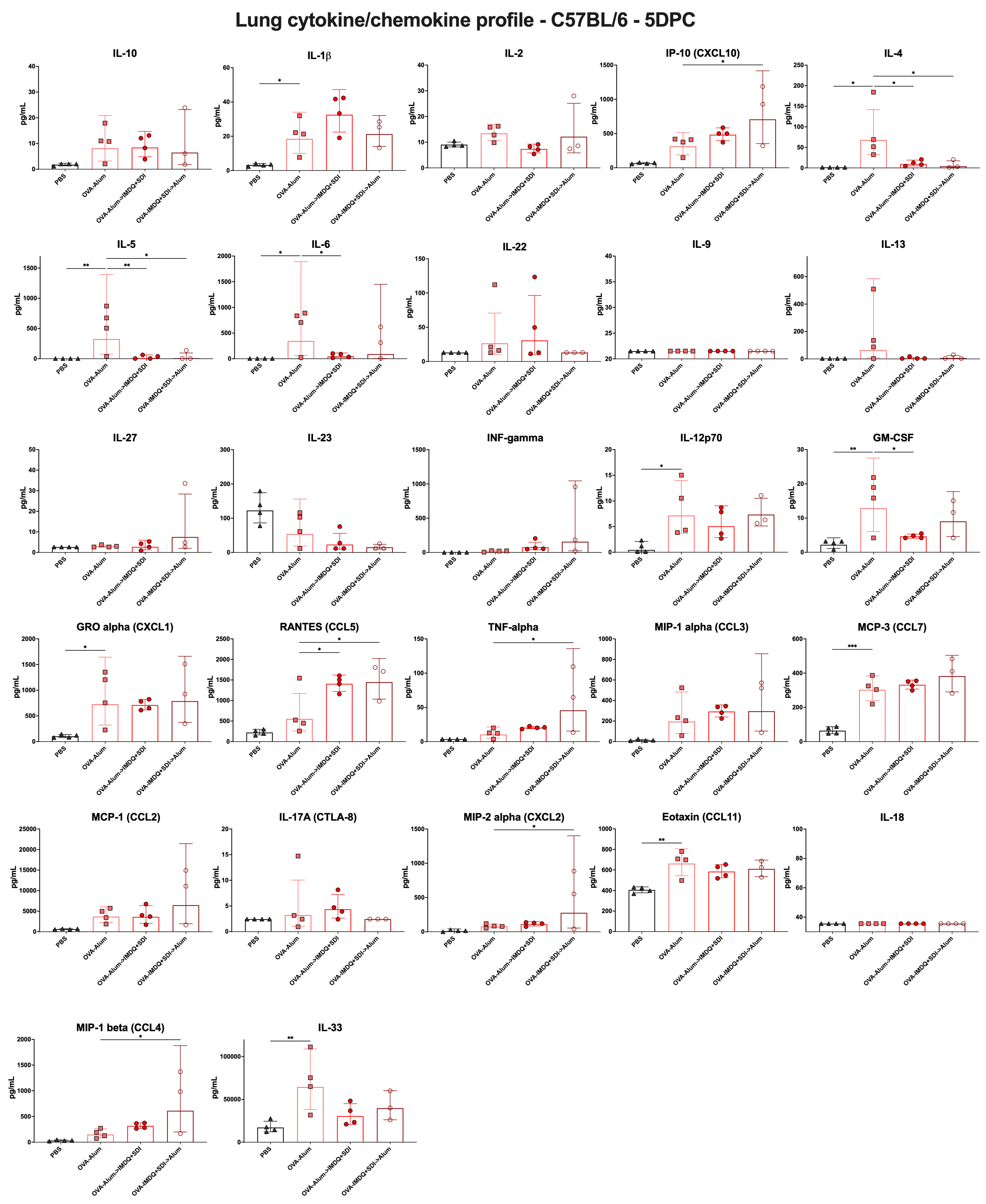

**Supplementary Fig. 4. Lung cytokine/chemokine profile in C57BL/6 primed with Alum or IMDQ+SDI and boosted with the other adjuvant.**

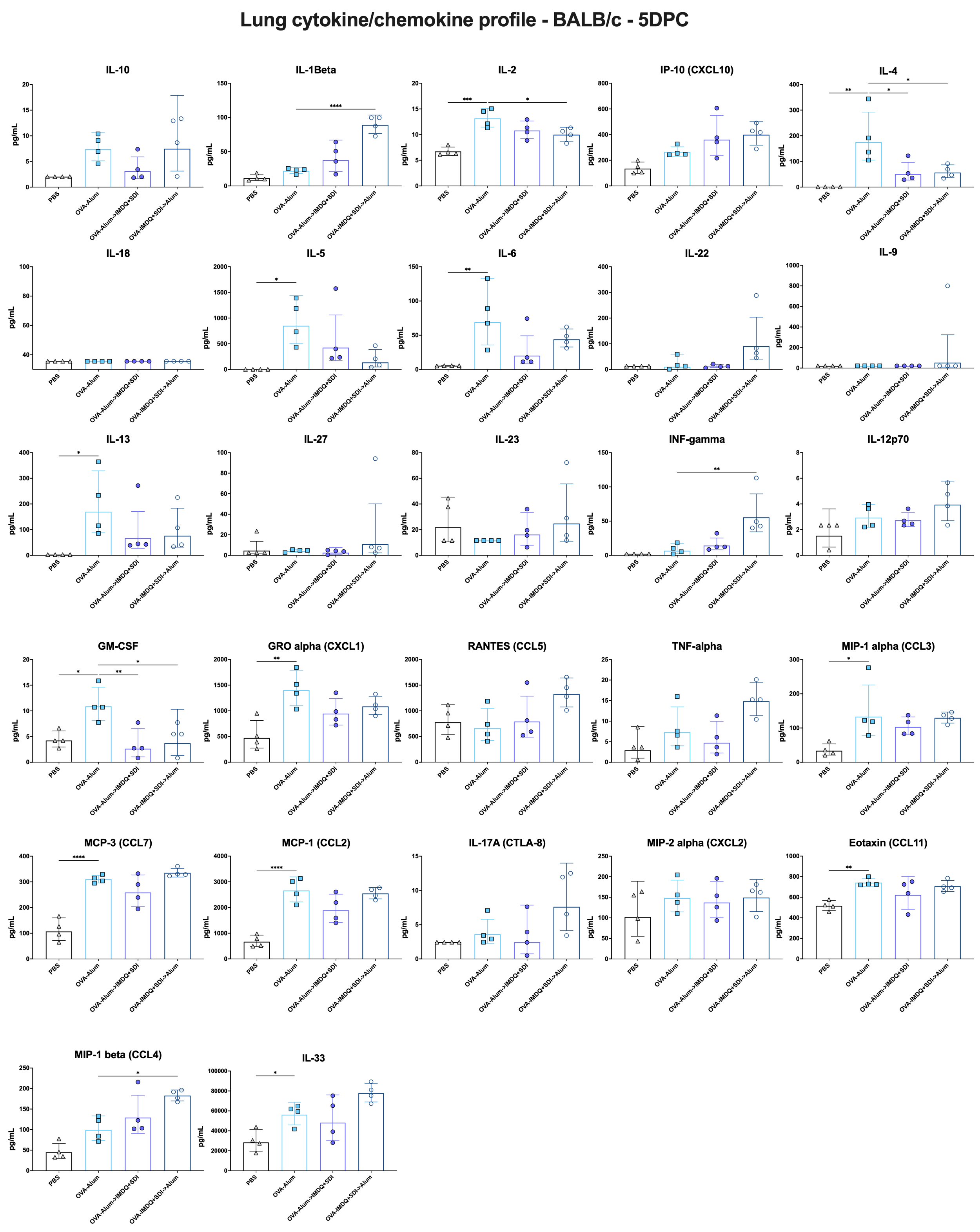

**Supplementary Fig. 5. Lung cytokine/chemokine profile in BALB/c primed with Alum or IMDQ+SDI and boosted with the other adjuvant.**

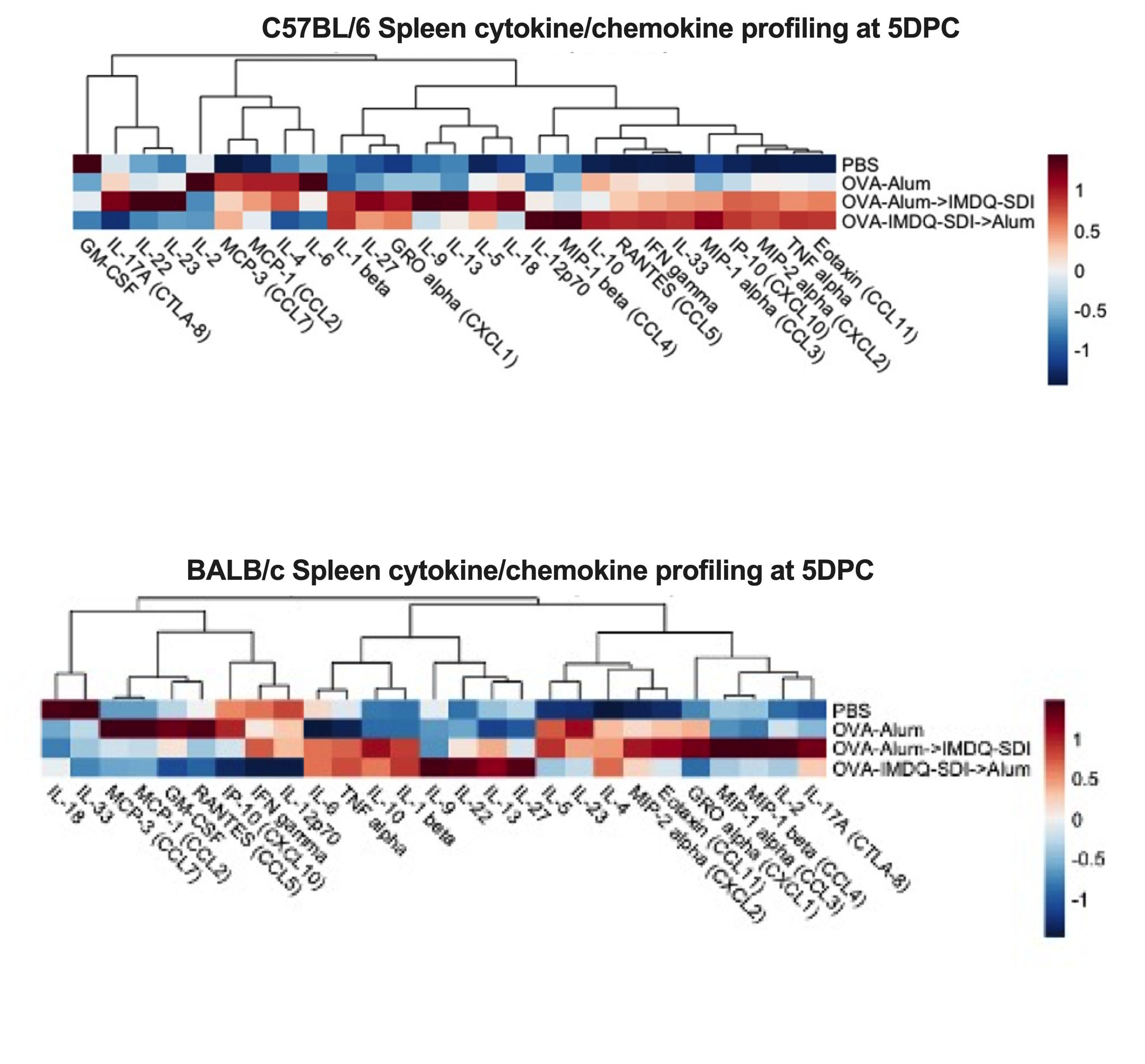

**Supplementary Fig. 6. Spleen cytokine/chemokine profile in C57BL/6 and BALB/c primed with Alum or IMDQ+SDI and boosted with the other adjuvant.**

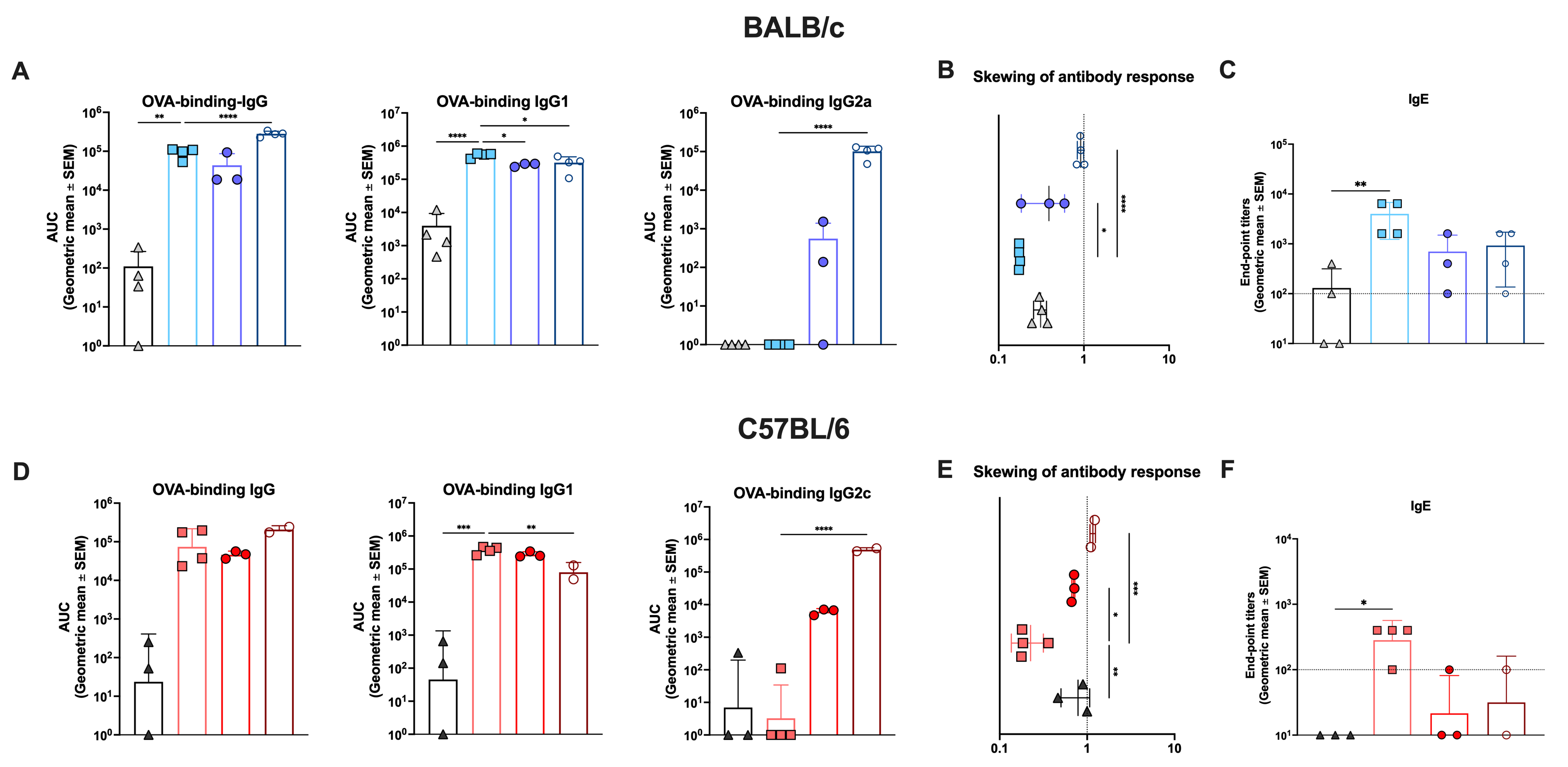

**Supplementary Fig. 7. OVA-specific and IgE antibody response after influenza challenge.**

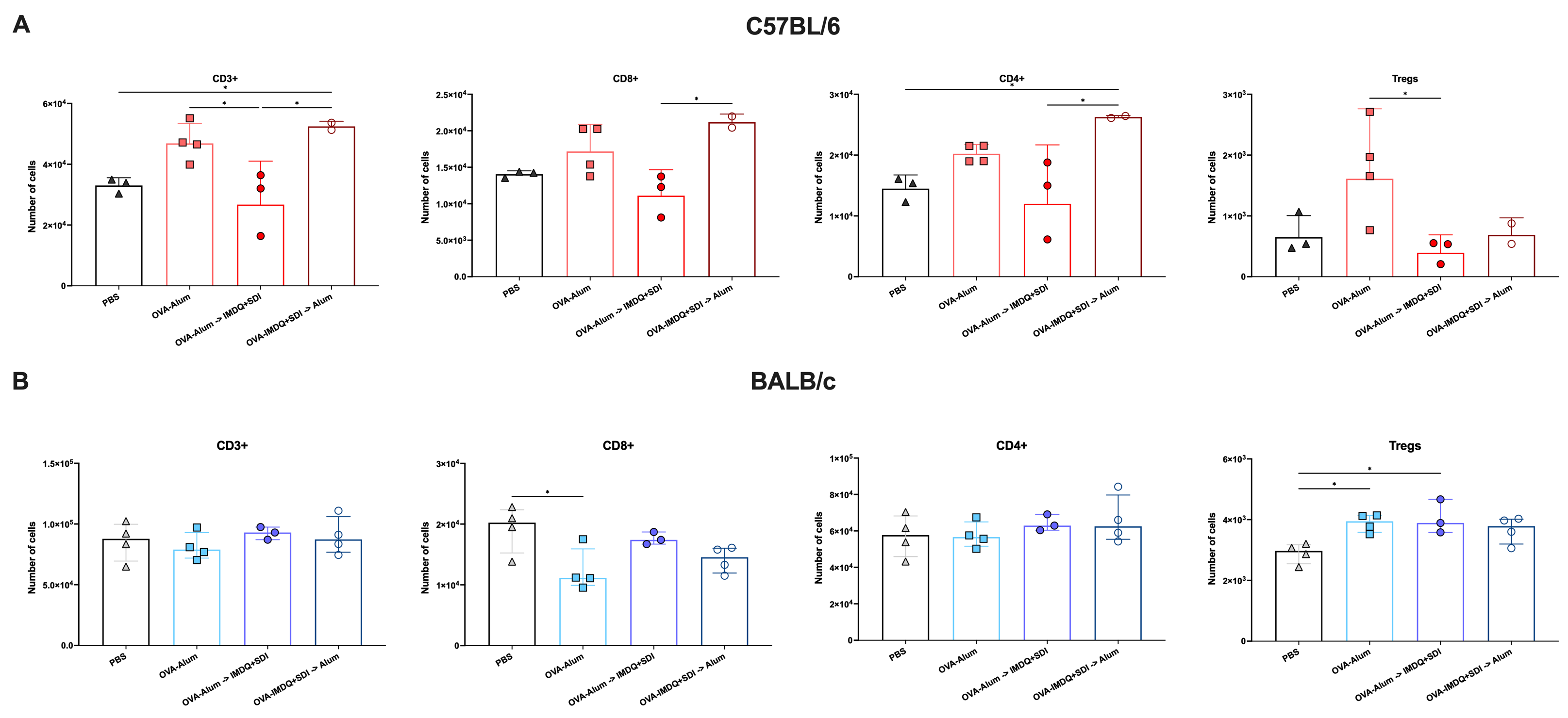

**Supplementary Fig 7. Number of cells in the lung for different T-cell subsets at 5DPI after influenza challenge.**

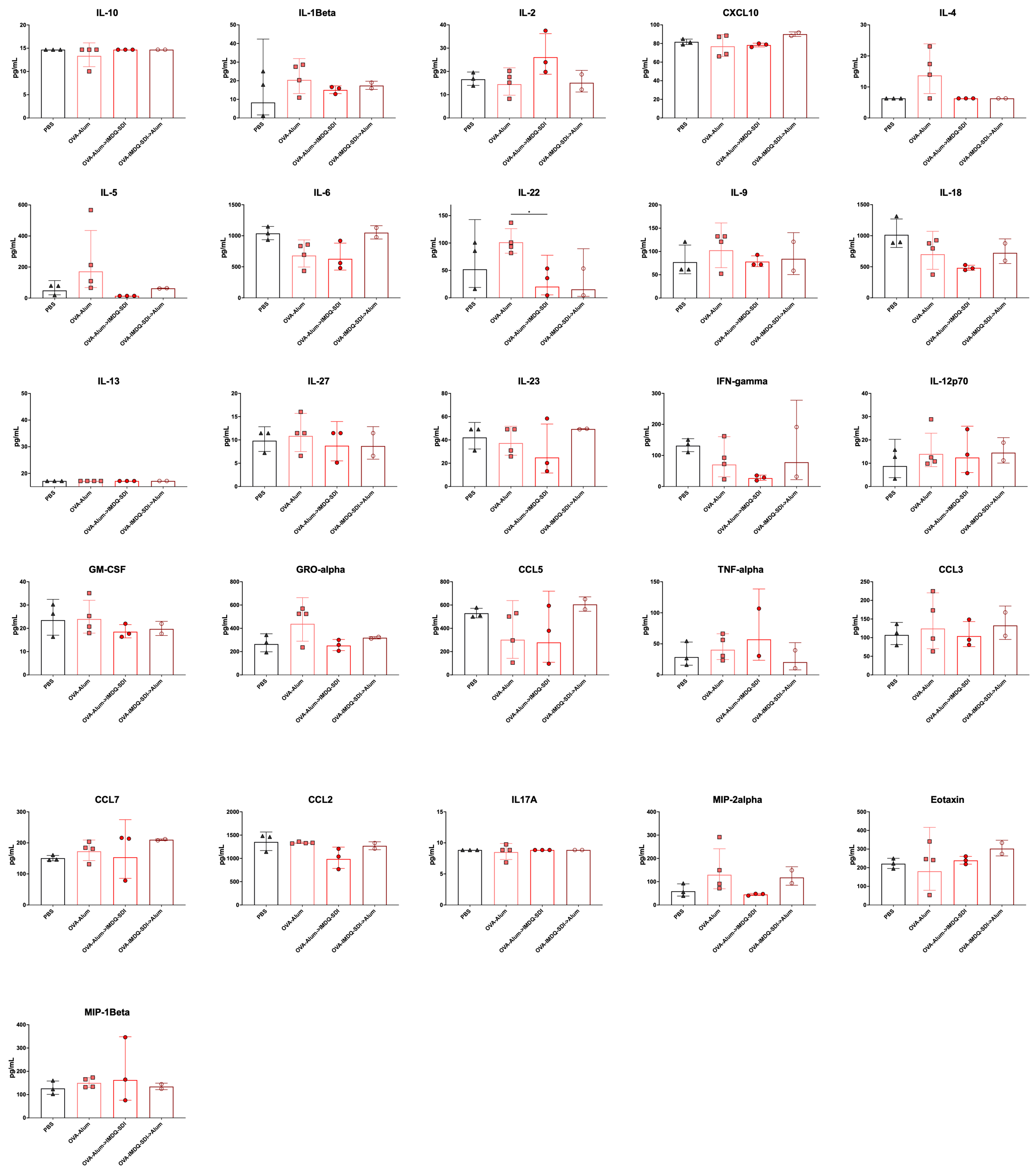

**Supplementary Figure 9. Lung Cytokine/chemokine protein concentration at 5DPI after influenza challenge in C57BL/6**

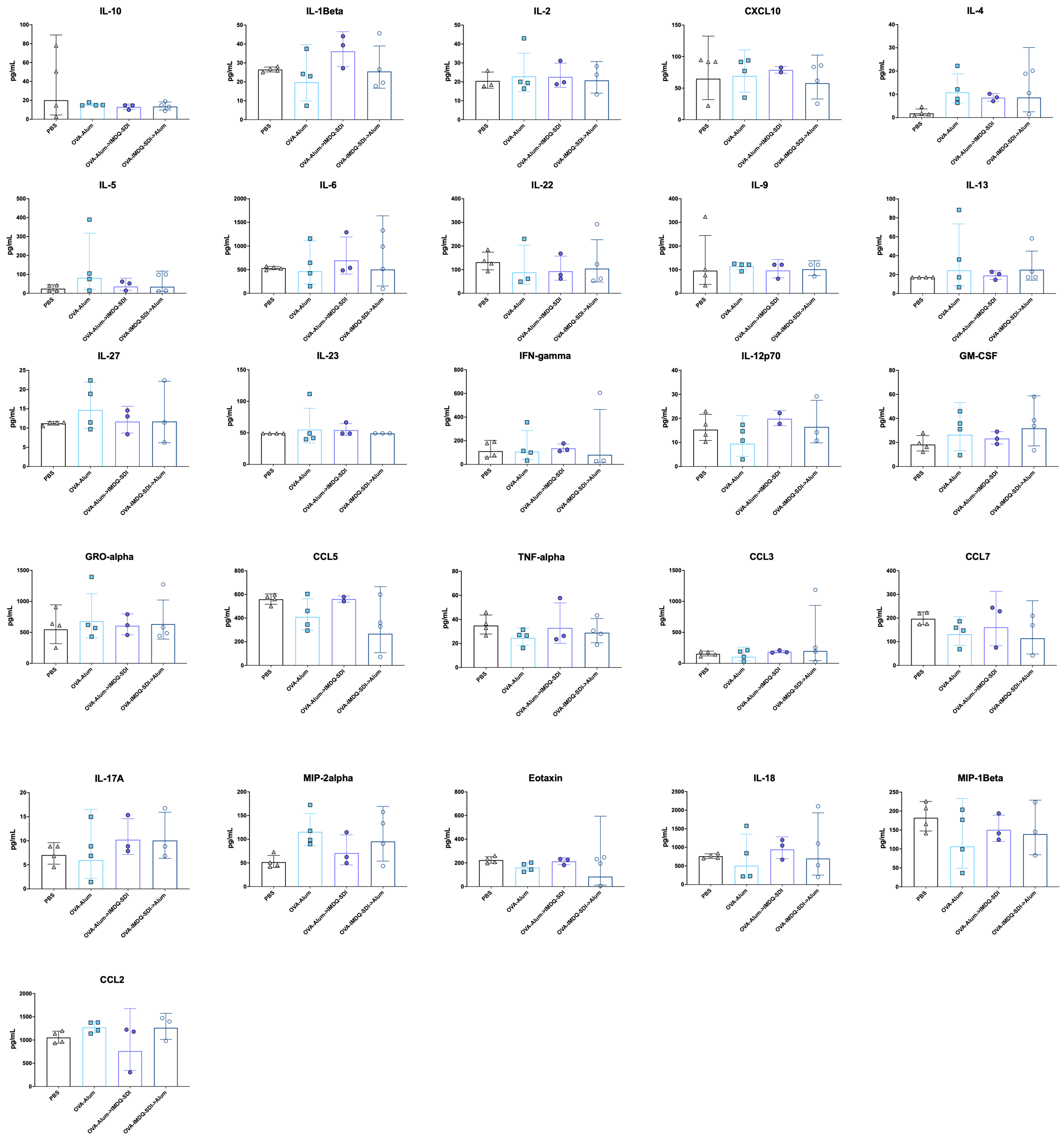

**Supplementary Figure 10. Lung Cytokine/chemokine protein concentration at 5DPI after influenza challenge in BALB/c**

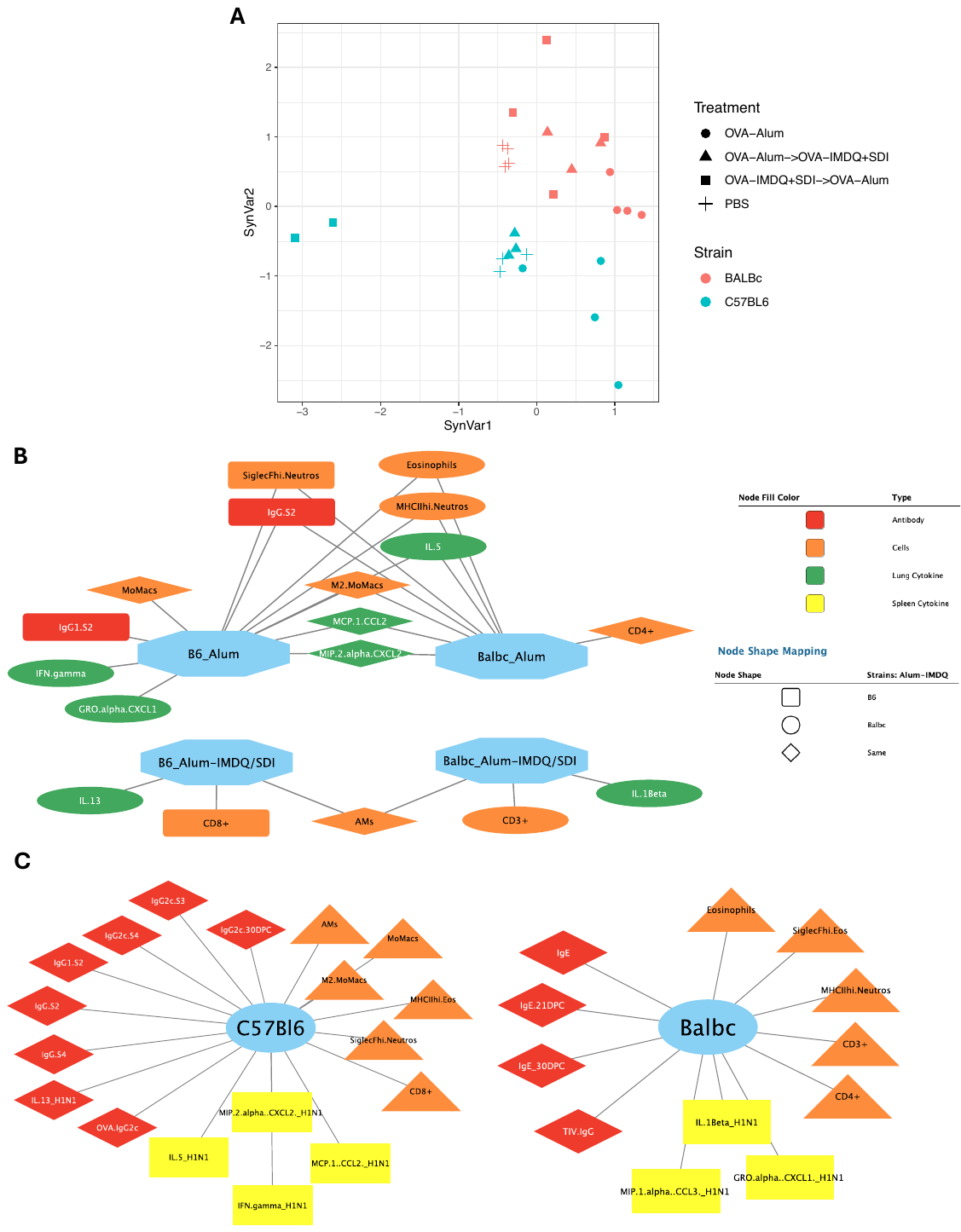

**Supplementary Figure 11. Multiple co-inertia analysis reveals similar effects on NC99 challenge outcome of OVA-sensitization in combination IMDQ+SDI in BALB/c and C57BL/6.**

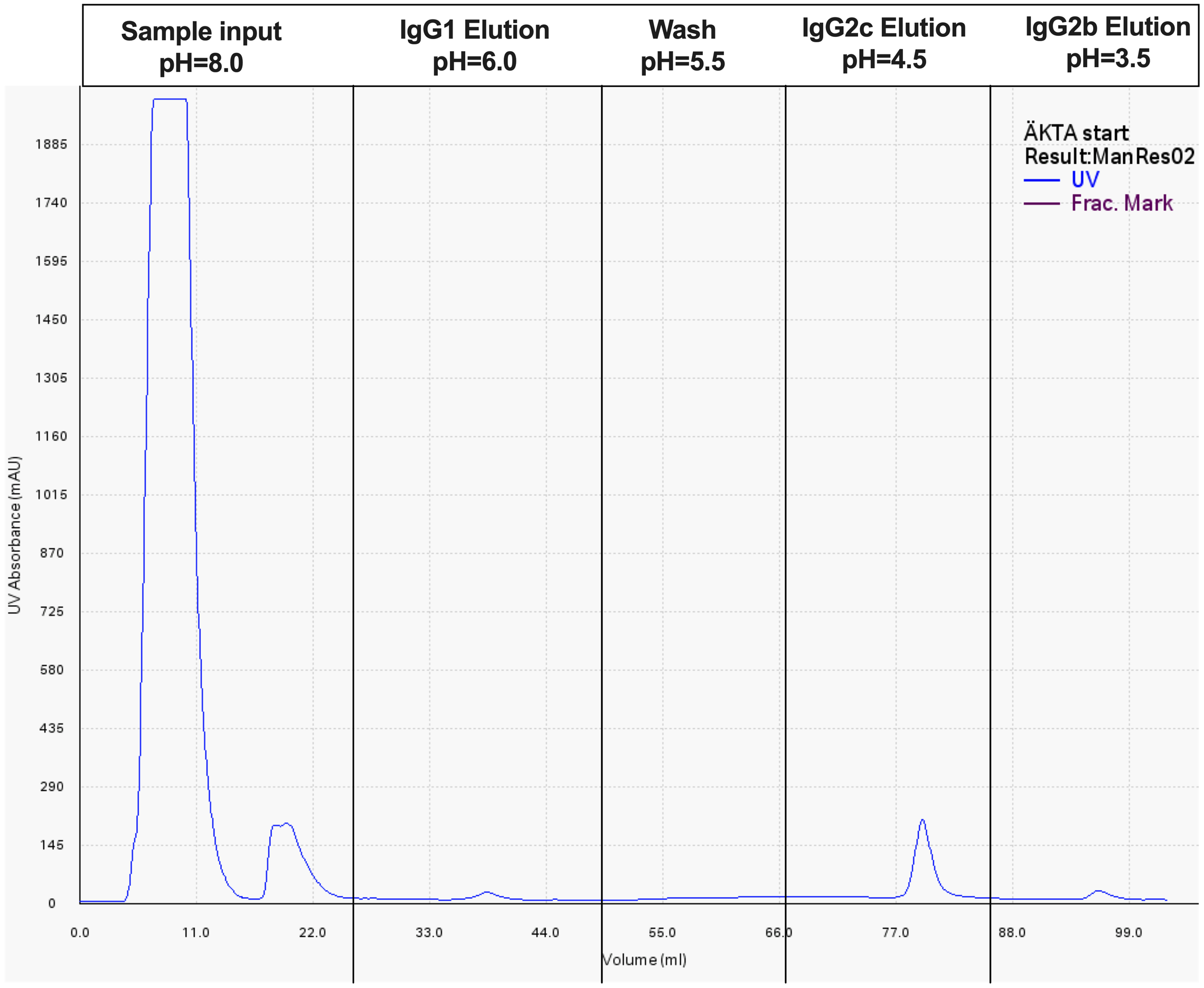

**Supplementary Figure 12. Purification of C57BL/6 IgG subclasses from serum using a Protein-A-sepharose column.**

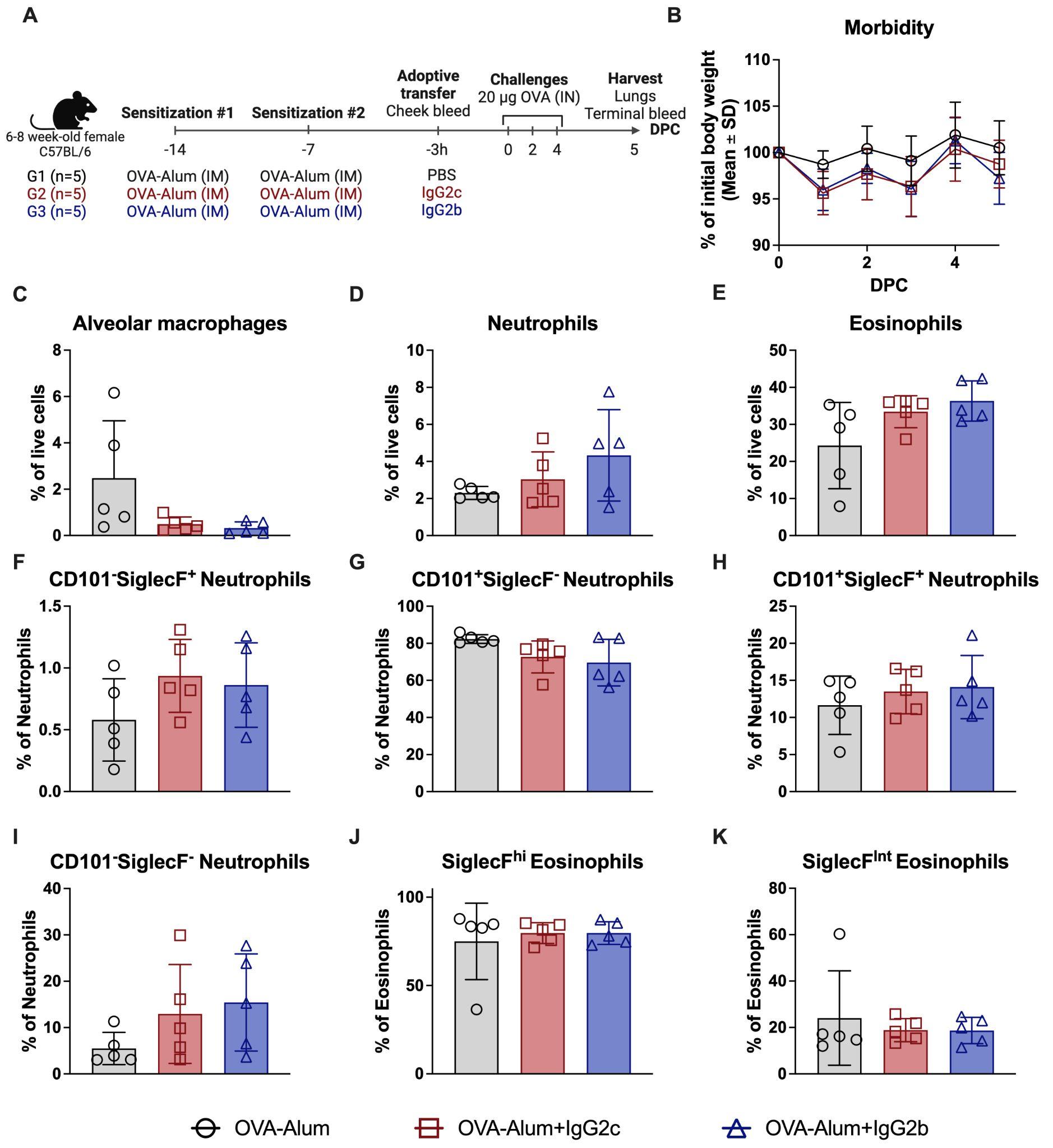

**Supplementary Figure 13. Transfer of IgG2c or IgG2b has no protective effect on in Alum-sensitized mice.** A) Experimental design. Created with: [Biorender.com](http://biorender.com). B) Daily mice bodyweights measured from 0DPC to 5DPC. Lung flow cytometry percentage of cell subpopulations at 5DPC (C-K). C) Alveolar macrophages (Gating: Live>TER-119^-^>Ly6G^-^>CD11c^+^SiglecF^+^>MHCII^+^). D) Total neutrophils (Gating: Live>TER-119^-^>Ly6G^+^>IL-5Ra^+^CD11b^+^). E) Total eosinophils (Gating: Live>TER-119^-^>Ly6G^-^>CD11b^+^> IL-5Ra^+^SiglecF^+^). F) CD101^-^SiglecF^+^ Neutrophils (Gating: Live> TER-119^-^>Ly6G^+^>IL-5Ra^+^CD11b^+^>CD101^-^SiglecF^+^). G) CD101^+^SiglecF^-^ Neutrophils (Gating: Live> TER-119^-^>Ly6G^+^>IL-5Ra^+^CD11b^+^>CD101^+^SiglecF^-^). H) CD101^+^SiglecF^+^ Neutrophils (Gating: Live> TER-119^-^> Ly6G^+^> IL-5Ra^+^CD11b^+^> CD101^+^SiglecF^+^). I) CD101^-^SiglecF^-^ Neutrophils (Gating: Live>TER-119^-^>Ly6G^+^>IL-5Ra^+^CD11b^+^>CD101^-^SiglecF^-^). J) SiglecF^hi^ Eosinophils (Gating: Live>TER-119^-^>Ly6G^-^>CD11b^+^> IL-5Ra^+^SiglecF^+^>CD101^+^SiglecF^hi^). K) SiglecF^int^ Eosinophils (Gating: Live>TER-119^-^>Ly6G^-^>CD11b^+^> IL-5Ra^+^SiglecF^+^>CD101^-^SiglecF^int^). Statistical analysis: for comparisons among groups at several time points: Two-way ANOVA with Tukey’s multiple comparisons test using the OVA-Alum (IP) group as control for multiple comparisons. Color of the significance markers indicates the comparison they represent; for comparisons among groups at a specific time point: Kruskal-Wallis one-way ANOVA with Dunn’s multiple comparisons test using the OVA-Alum (IP) group as control for multiple comparisons.
